# Gut *Eggerthella lenta* promotes the efficacy of resveratrol through reductive metabolism

**DOI:** 10.1101/2024.03.10.584274

**Authors:** Zhixiang Dong, Peijun Yu, Hui Zhou, Rui Li, Qiang Sun, Yunpeng Yang, Yang Gu, Weihong Jiang

## Abstract

Resveratrol (RSV) is a plant-derived natural product with diverse biological activities. It has attracted considerable attention for its notable efficacy in nutritional health and disease treatment. The physiological impact of RSV in the human body is closely connected to the gut microbiota; however, the primary gut microorganisms responsible for RSV metabolism and the underlying mechanisms remain unclear. Here, based on the *ex vivo* culturing of the gut microbiota from human feces, we isolated a bacterium capable of efficiently metabolizing RSV, namely *Eggerthella lenta* J01. Through the induced enrichment transcriptomics and bioinformatic analyses, we further identified a resveratrol reductase (RER) from *E. lenta* J01, which specifically catalyzes the hydrogenation of the C9–C10 double bond of RSV and initiates RSV’s in vivo metabolism. RER and its homologs represent a novel class of ene-reductases. The abundance of RER in the gut microbiota of healthy individuals was significantly higher than that in patients with inflammatory bowel disease, suggesting its crucial physiological function. Cell culture experiments and animal studies showed that dihydroresveratrol (DHR), a metabolite of RSV catalyzed by RER, exhibited stronger biological activity in inhibiting the growth of colon cancer cells and alleviating symptoms of enteritis in mouse models. Our results expand the understanding of gut microbial metabolism of RSV and its medicinal functions, providing possible guidance for optimizing RSV bioavailability in the human body.

## Introduction

Plant-derived foods are rich in natural bioactive compounds, which are intricately linked to human health. In recent years, a large number of studies have demonstrated a significant impact of the gut microbiota on the in vivo function of plant-derived compounds^1^. The gut microbiota harbors a comprehensive enzymatic system, and thus, these microorganisms can metabolize a large number of natural compounds that we ingest, e.g., plant lignans^2^, orange fiber^1^, and cardiac drug digoxin^3^, and modify their activity and bioavailability. Such an influence includes either deactivation or activation of plant-derived natural products, which changes their physiological functions^4^.

Resveratrol (RSV) is a natural polyphenol abundant in various berries and foods^5^. It has diverse biological activities such as antitumor^6^, anti-inflammatory^7^, anti– cardiovascular disease^8^, and antioxidant effects^9^. RSV exhibits a significant effect on gut microbiota. Specifically, following RSV intake in mice, a notable modification occurs in the composition of gut microbiota, with an increase and decrease in the relative abundance of some beneficial and pathogenic bacteria, respectively^10^. The regulatory effect of RSV on the proportion of beneficial bacteria in intestinal microbiota inhibits the formation of the harmful trimethylamine (TMA), thereby reducing the incidence of cardiovascular diseases in mice^8^. Furthermore, RSV has a role in maintaining the homeostasis of lipid metabolism by selectively modulating gut microbiota ecology^11^.

RSV is metabolized by gut microbiota, and it has been shown that RSV can be converted into multiple small-molecule compounds by gut microbiota^12^, and many of these metabolites exhibit various medicinal effects^13^. These findings suggest the importance of gut microbiota in assisting RSV to exert its efficacy in the human body. Nevertheless, it is still largely unknown which gut microorganisms metabolize RSV, and the specific mechanism remains unclear. To date, very few intestinal strains (i.e., *Slackia equolifaciens* and *Adlercreutzia equolifaciens*) have been found to be able to metabolize RSV, but their metabolic capacity for RSV is poor^12^. The major strains as well as the key enzymes responsible for RSV metabolism in the human body remain unknown.

Here, using a combination of enrichment culturing and 16S rRNA amplicon sequencing, we discovered a gut bacterial strain *Eggerthella lenta* J01 from fecal samples of healthy individuals, which is capable of efficiently metabolizing RSV. Through further comparative transcriptomics and biochemical analyses, we identified and functionally characterized a novel RSV reductase (RER), capable of efficiently catalyzing the reduction of RSV toward dihydroresveratrol (DHR) in *E. lenta* J01. Interestingly, RER and its homologs represent a novel class of ene-reductases. The presence of RER significantly enhanced the inhibitory effect of RSV on colorectal cancer cells. Furthermore, in a colitis mouse model, we showed that the efficacy of RSV in alleviating colitis depended on its RER-mediated conversion into DHR. Of note, the abundance of RER-encoding genes in the healthy human gut microbiota is much higher than that in colitis patients, suggesting the importance of RERs in RSV-related protection against colitis.

## Results

### Discovery of the intestinal microorganisms capable of efficiently metabolizing RSV

After ingestion, RSV can be metabolized by gut microbiota to form small-molecule metabolites with various physiological activities^12,13^ (Fig. 1A). To determine whether there are potential key intestinal strains in gut microbiota responsible for RSV metabolism, we collected fecal samples from three healthy volunteers for *ex vivo* cultivation and then assessed the difference in RSV metabolism between gut microbiota and *S. equolifaciens*, a reported gut bacterium with weak metabolic capacity to RSV. We found that the collected gut microbiota metabolized RSV at the rate of 7 μM/h, much higher than the rate of *S. equolifaciens* (Fig. 1B), suggesting the existence of major intestinal strains capable of metabolizing RSV with high efficiency.

**Fig. 1.**
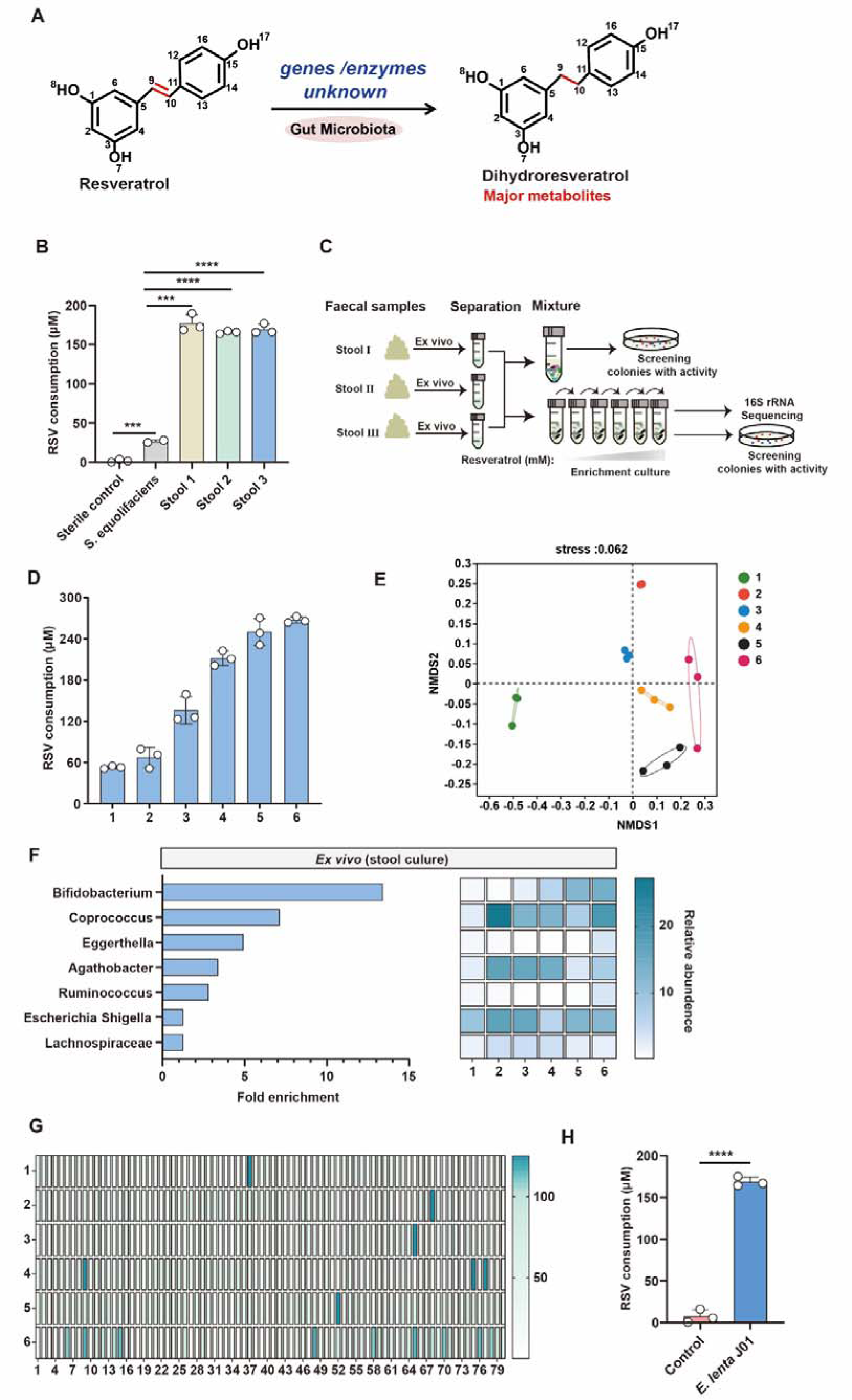
| Discovery of resveratrol (RSV)-metabolizing *E. lenta* J01 strain from gut microbiota. **A**, Proposed microbial metabolic reaction for RSV reduction. **B**, RSV consumption by the microbiota from three fecal samples and by *S. equolifaciens* (a reported gut bacterium capable of metabolizing RSV). Statistical analysis was performed by a two-tailed Student’s *t*-test (***, p<0.001; ****, p<0.0001). **C**, Schematic presentation of isolating efficient RSV-consuming gut strains from human fecal samples. **D**, Gradually increased RSV consumption by gut microbiota via enrichment culturing in the presence of RSV. Data are represented as mean ± s.d. (*n* = 3). Error bars show s.d. **E**, The changes in gut microbial composition and structure after different rounds of enrichment culturing based on the NMDS analysis. The dots and annuli with different colors represent samples of different groups. The distance between two dots represents the similarity of the species composition of the corresponding samples. Shorter distance indicates higher similarity. **F**, Identification of the taxonomic classification of the RSV-metabolizing microorganisms (generated from enrichment culturing) via 16S rRNA amplicon sequencing. Fold enrichment represents the increase of these gut strains after enrichment culturing. The heat map represents the relative abundance of each strain in the gut microbiota after different rounds of enrichment culturing. **G**, Isolation of the gut strain capable of consuming RSV. The isolates for testing were cultivated anaerobically in the mGAM medium supplemented with RSV (150 μM) for 24 h. Among the 480 tested isolates, 20 were found to be able to efficiently metabolize RSV, and all of these strains were identified as *E. lenta* strains. **H**, Evaluation of the RSV consumption by the *E. lenta* J01 strain. Data are presented as mean ± s.d. (*n* = 3). Error bars show s.d. Statistical analysis was performed by a two-tailed Student’s *t*-test (****, p<0.0001).

To identify these intestinal strains, we cultured the aforementioned gut microbiota samples using the previously reported *ex vivo* system in the presence of RSV, aiming to enrich gut microorganisms that have high capability to metabolize RSV (Fig. 1C). We found that, with the increase of enrichment rounds, the ability of intestinal microbiota to metabolize RSV gradually improved, increasing from 60 μM RSV consumption every 24 h for the first-generation culture to 240 μM RSV consumption every 24 h for the sixth-generation culture (Fig. 1D). Next, we performed 16S rRNA amplicon sequencing analysis on the 18 enriched cultures (three fecal samples for six rounds of enrichment) (Fig. 1C). The results showed that the stepwise enrichment with RSV significantly altered the composition of gut microbiota (Fig. 1E-F) (Table S1) in which the abundance of Actinobacteria, particularly *Bifidobacterium* and *Eggerthella*, was greatly enhanced (Fig. S1).

Based on these findings, we sought to further isolate Actinobacteria strains with high RSV-metabolizing ability from the abovementioned enriched cultures. To this end, one aliquot of culture was plated on the solid BHI medium (suitable for cultivating Actinobacteria) to obtain individual colonies, and then a total of 480 isolates were randomly selected to examine their ability in anaerobic RSV metabolism. Twenty isolates exhibited a high capacity for RSV metabolism (Fig. 1G and Fig. S2), with the highest RSV-consumption rate of ∼16.7 μM/h (200 μM within 12 h) (Fig. 1H). Further identification using 16S rDNA full-length sequencing indicated that all these 20 isolates belong to *Eggerthella lenta*. Next, we chose the isolate with the highest RSV-metabolizing capability for genomic sequencing (accession number PRJNA1076688), and found that the genome size of this strain was 3,311,593 bp (Fig. S3), smaller than the genome size of all reported *E. lenta* strains (Table S2). Here, we termed this strain *E. lenta* J01. The phylogenetic analysis showed that *E. lenta* J01 has a distant evolutionary relationship with other *E. lenta* strains (from the NCBI database) (Fig. S4), suggesting that it is an unreported *E. lenta* strain.

### Identification of key enzymes in RSV metabolism

The above findings support that *E. lenta* J01 plays an important role in converting RSV. Next, we attempted to identify the key enzyme(s) responsible for this metabolic process in *E. lenta* J01. To this end, we adopted an RSV-induced transcriptional profile to find potentially essential genes for RSV consumption. In brief, *E. lenta* J01 was cultivated in the mGAM medium with or without the addition of RSV, and then subjected to a comparative transcriptomic analysis using RNA sequencing (Fig. 2A). Compared with the sample without RSV, the addition of RSV to the medium led to significant transcriptional changes (|log fold-change| ≥1) of 491 genes, where 283 genes were upregulated (Fig. 2B) (Table S3). Considering that the first step of in vivo RSV metabolism has been supposed to be a reduction reaction to form DHR (Ref), we specifically focused on the greatly upregulated genes (|log fold-change| ≥2) that encode redox-related enzymes and selected 21 genes for the test (Table S4). These genes were then cloned and expressed separately in *E. coli* BL21, and the resulting recombinant strains were used to examine the activity for RSV metabolism. Encouragingly, efficient RSV consumption was found in the *E. coli* strain expressing the LUA64_RS01375 gene (Figs. 2C–D and Fig. S5), whereas no obvious changes occurred in the other *E. coli* strains. Noticeably, the LUA64_RS01375 gene exhibited the highest upregulation in the presence of RSV according to the abovementioned RNA-sequencing data (Table S3). Therefore, these findings strongly suggest that the protein encoded by LUA64_RS01375 is the target enzyme. We named this enzyme RSV reductase (RER).

**Fig. 2.**
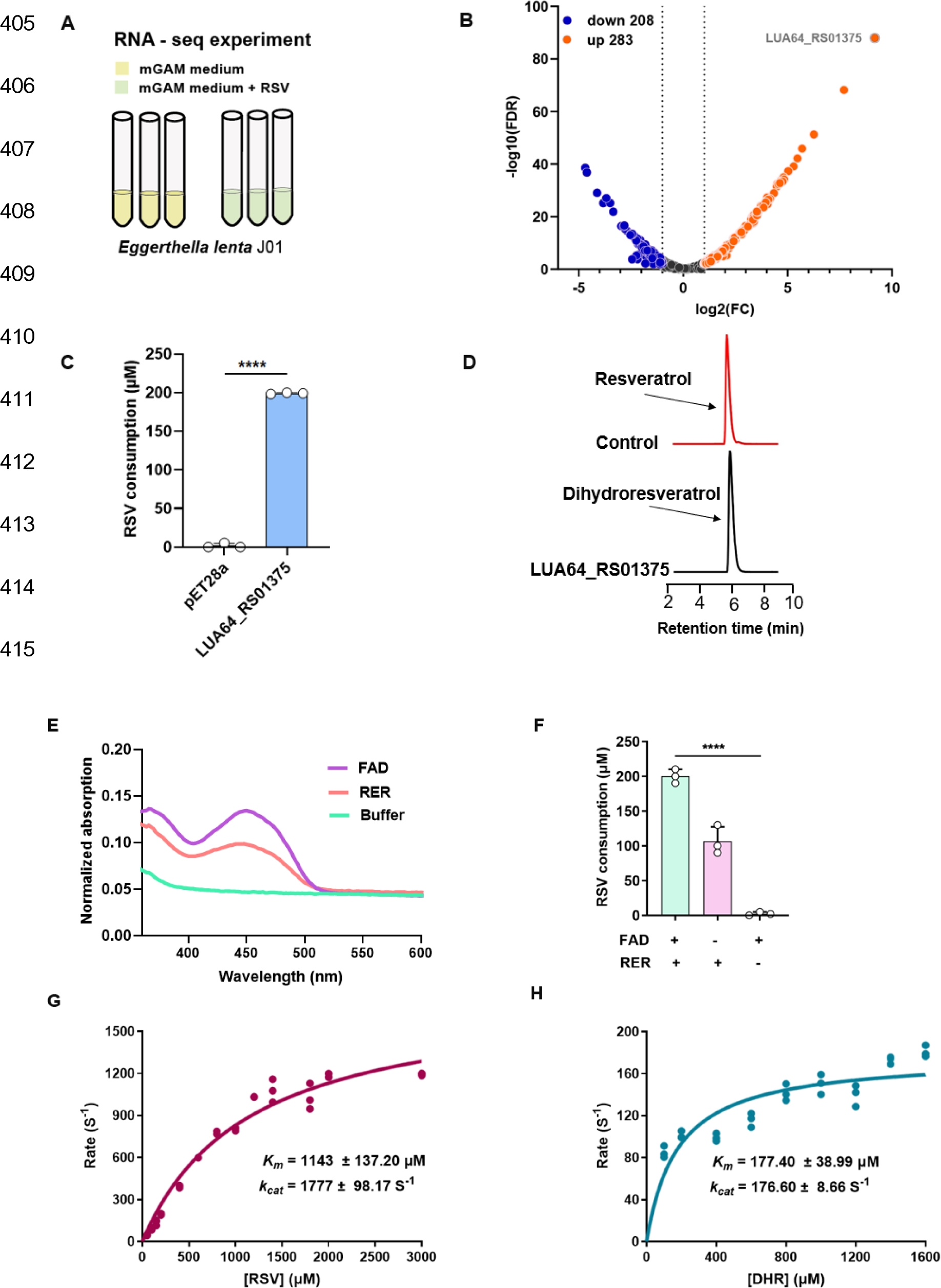
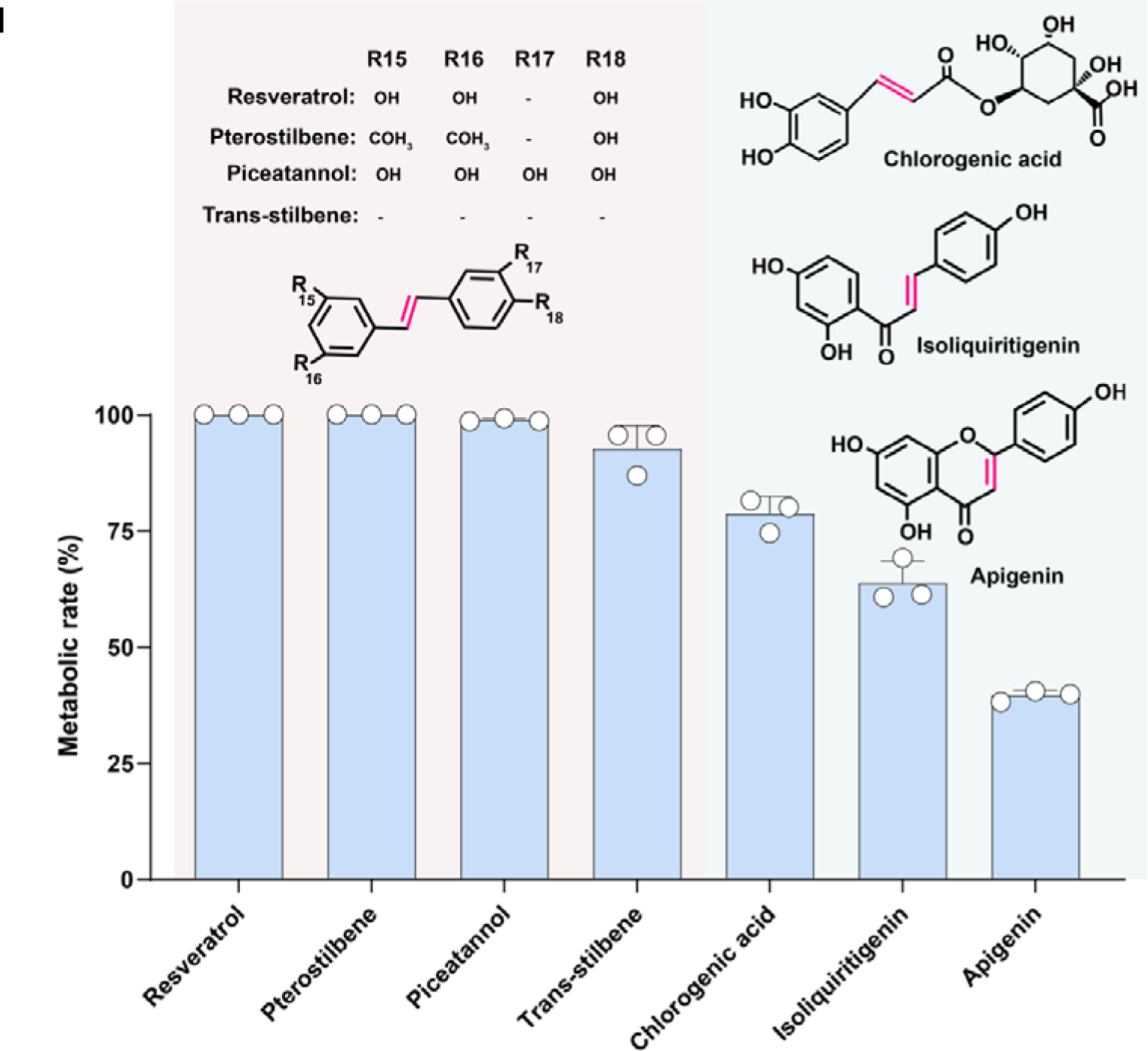
| Discovery of RSV-metabolizing enzyme in *E. lenta* J01. **A**, Design of an RSV-induced culturing system for RNA sequencing to identify potential genes that encode RSV-metabolizing enzymes in *E. lenta* J01. The presence of RSV in the medium was expected to cause the upregulation of target genes. **B**, Volcano map of the differentially expressed genes in *E. lenta* J01 in the presence (100 μM) and absence of RSV. Each dot represents one gene. The red and blue dots indicate the genes with significant upregulation and downregulation, respectively. The other genes are represented by grey dots. FC, fold change. **C, D**, HPLC detection of the conversion of RSV into DHR by the *E. coli* strain (BL21) expressing the LUA64_RS01375 gene. Data are represented as mean ± s.d. (*n* = 3). Error bars show s.d. Statistical analysis was performed by a two-tailed Student’s *t*-test (****, p<0.0001). **E**, Ultraviolet-visible absorption spectroscopy of FAD (standard) and the purified RER protein. The same characteristic absorption peaks were observed for both RER (red) and FAD (purple). **F**, In vitro validation of the RSV-metabolizing activity of the purified RER protein. Data are represented as mean ± s.d. (*n* = 3). Statistical analysis was performed by one-way annova (****, p<0.0001). **G, H**, In vitro steady-state kinetic analysis of RER using RSV or DHR as the substrate. Each data point represents mean ± s.d. (*n* = 3). Error bars show s.d. **I**, A screening of the substrate spectrum of RER. Initial turnover rates of the purified RER protein toward different substrates (100 μM) were determined. Data are represented as mean ± s.d. (*n* = 3).

To further elucidate the catalytic property of RER, we expressed and purified it for in vitro assays. The analysis of the visible absorption spectra of RER revealed a characteristic absorption peak (450 nm), which was the same as that of the flavin cofactor, flavin adenine dinucleotide (FAD) (Fig. 2E). Therefore, a reaction system coupling methyl viologen consumption and FADH_2_ production was built to analyze the RER activity toward RSV. Expectedly, the reduction of RSV and the formation of DHR were detected in the presence of RER (Fig. 2F and Fig. S6). Afterward, we determined the optimal reaction conditions (pH 7.5 and temperature 37°C) for RER (Fig. S7). Under these conditions, RER gave a *K*_m_ of 1143 ± 137.2 μM and a *k*_cat_ of 1777 ± 98.17 s^−1^ toward RSV. In addition, we found that RER was able to catalyze the conversion of DHR into RSV with a lower *k*_cat_/*K*_m_ value (Figs. 2G–H).

To determine the substrate range and preference of RER, we selected a series of natural dietary and medicinal compounds with a similar structure to RSV for testing. We found that RER had catalytic activity toward trans-stilbene, pterostilbene, piceatannol, apigenin, isoliquiritigenin, and chlorogenic acid (Fig. 2I), but no definite activity toward cis-resveratrol, naringenin, and p-hydroxycinnamic acid (Fig. S8).

### The RER-like enzymes represent a distinct class of ene-reductases

Next, we investigated the distribution of RER-like enzymes in gut microbiota using the amino acid sequence of RER in *E. lenta* J01 as the query. Based on a BLASTP search, we identified 63 putative RER enzymes (amino acid sequence identity ≥59%, coverage ≥56%, e-value ≤1e−10), covering some uncategorized species, for a maximum-likelihood phylogenetic analysis (Fig. 3A). The results revealed that most of the RER enzymes exist in five genera of gut bacteria, namely *Eggerthella*, *Gordonibacter*, *Slackia*, *Raoultibacter*, and *Adlercreutzia*, and the genus *Eggerthella* occupied the largest proportion (Fig. 3A). All of the abovementioned bacteria belong to phylum Actinobacteria (Fig. 3A). To further validate the reliability of the phylogenetic analysis, we selected four putative RER enzymes to assess their activity toward RSV. As expected, all of the enzymes were able to catalyze the transformation of RSV to DHR, although with different catalytic efficiency (Fig. 3A).

**Fig. 3.**
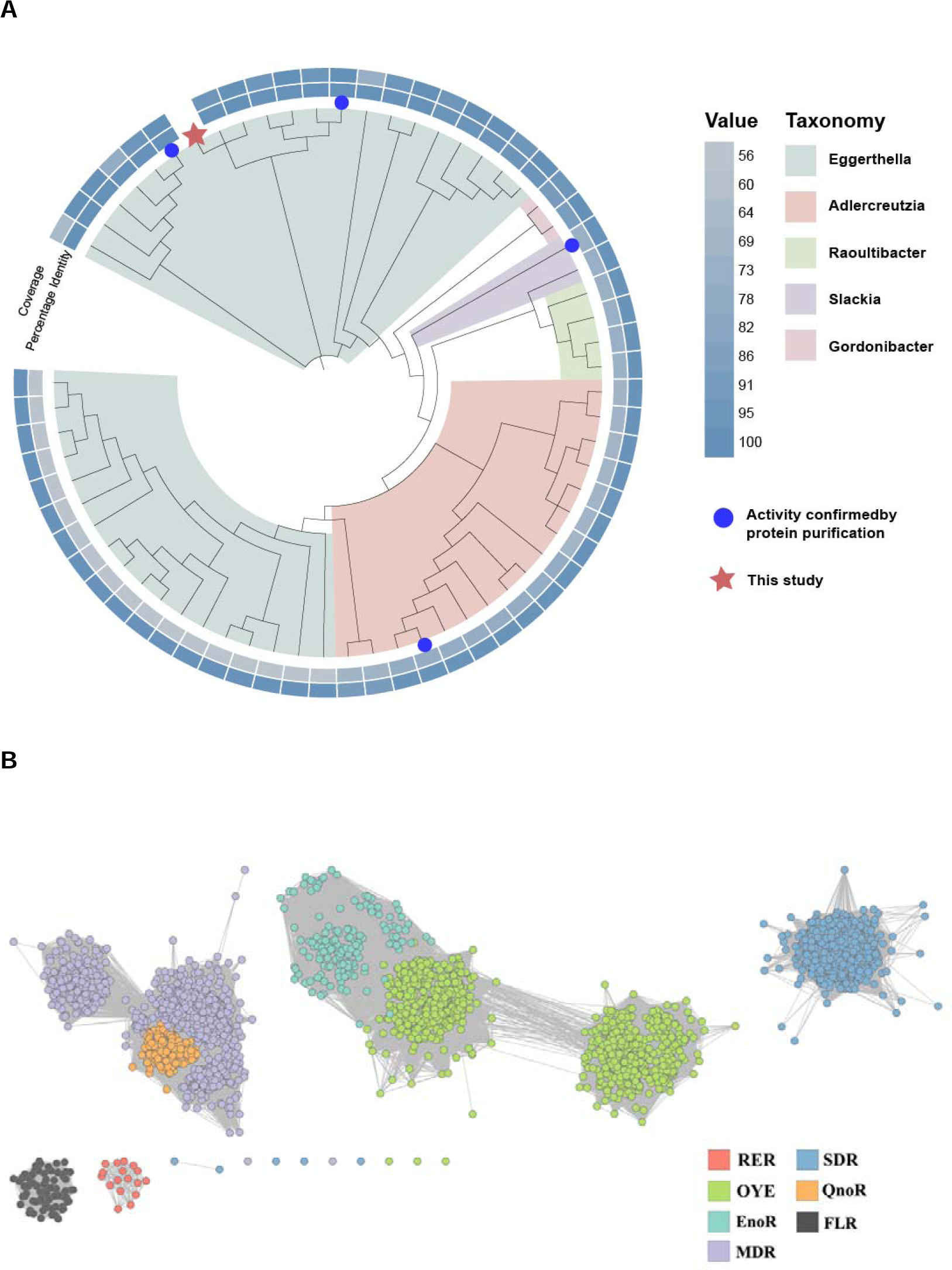
| Identification of resveratrol reductases (RERs) as a distinct group of ene-reductases based on bioinformatic and experimental analysis. **A**, Phylogenetic analysis of the RER enzymes. The maximum-likelihood phylogenetic tree is based on the protein sequences of the putative RER enzymes found in *Eggerthella*, *Gordonibacter*, *Slackia*, *Raoultibacter*, and *Adlercreutzia*. The blue circles in Fig. 3A indicates that RER homologous proteins have been shown to have catalytic RSV-reduction activity in vitro. The depth of the outer ring color represents the coverage and the percentage identity of the RER homologous protein, where darker color indicates higher similarity. **B**, A sequence similarity network (SSN) for the ene-reductases. The network was generated with an initial score of 10^−20^. The “alignment score threshold” (a measure of the minimum sequence similarity threshold) for drawing the edges that connect the proteins (nodes) in the SSN was refined such that nodes are connected by an edge if the value is ≥40. Each of the nodes represents the proteins with ≥95% amino acid sequence identity and is colored based on the cluster types (OYE, EnoR, SDR, MDR, QnoR, FLR, and RER). Each type of ene-reductase (highlighted with different colors) is separated into different clusters that may include enzymes with similar biochemical activity. A total of 63 different RER-like enzymes shown in the phylogenic tree are denoted in red. The nodes representing the reported ene-reductase with enzyme activity data (from the KEGG database) and identified RER enzymes in this work are highlighted with light red.

To date, the enzymes capable of catalyzing the asymmetric activation of olefin reduction are assigned as ene-reductases^14^. According to the catalytic property, the RER enzyme discovered in this study also belongs to ene-reductases; however, the result of BLASTP analysis showed that RER has a very low amino acid sequence similarity to currently reported ene-reductases (Table S5), indicating a remote evolutionary relationship between RER and those reductases. To confirm this, we constructed a protein sequence similarity network (SSN) using the aforementioned 63 RER homologous sequences and nearly 6818 sequences derived from six known ene-reductase families (based on the KEGG database) (Table S6). Consequently, RER-like enzymes represent a distinct class of ene-reductases (Fig. 3B).

### The RSV’s metabolite DHR exhibits stronger antitumor cell activity

Previous in vitro studies have demonstrated that both RSV and its metabolites have an inhibitory effect on the proliferation of colorectal cancer cells^15^. Here, we explored whether the *E. lenta* J01 strain and the RER enzyme are involved in this biological process. Hence, we coincubated the *E. lenta* J01 strain with RSV for 12 h, and then collected the supernatant to analyze RSV consumption and product formation. We found that RSV was completely converted into the expected product, DHR (Fig. 4A). Next, the culture medium was aseptically treated and cocultured with the colorectal cancer cell line HCT-116, with untreated RSV serving as the control group. Although even RSV itself was able to inhibit the proliferation of colorectal cancer cells (Fig. 4B), the reaction liquid containing its metabolite DHR had a more pronounced inhibitory effect on cell proliferation (> 30%) (Fig. 4B). Subsequently, we overexpressed RER in the *E. coli* BL21 strain, yielding the recombinant strain BL21 (RER) for further testing. The results from the cell intervention experiments showed that, compared with the control group treated with RSV alone, the cell-free reaction liquid coincubated with BL21 (RER) and RSV had a stronger inhibitory effect on the proliferation of HCT-116 cells (> 30%) (Fig. 4C). These findings demonstrate that the RER-mediated metabolic reaction in *E. lenta* J01 can generate RSV derivatives with stronger antitumor activity.

**Fig. 4.**
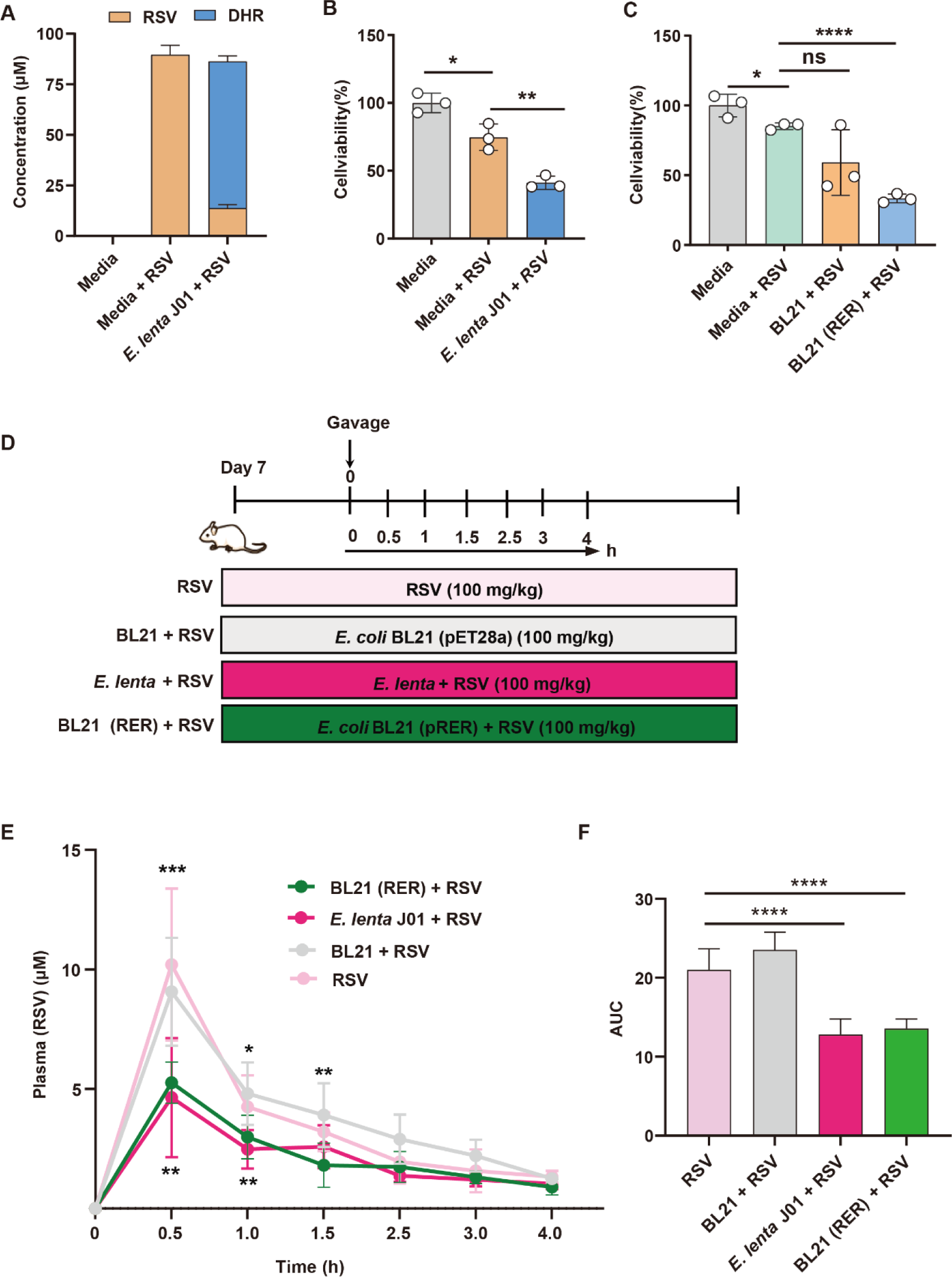
| Effects of RSV and its metabolite DHR on tumor cells and the pharmacokinetics of RSV in the blood of mice. **A**, HPLC detection of the predicted RSV metabolite DHR. The reaction was carried out anaerobically for 24 h (Media, Media + 100 μM RSV, *E. lenta* + 100 μM RSV). **B, C**, The effects of RSV and its metabolite DHR on tumor cells mediated by *E. lenta* (**B**) and *E. coli* BL21 (RER) (**C**). Colorectal cancer cell HT116 was cultured under different conditions (Media, Media + RSV, empty vector + RSV, and *E. coli* BL21_RER + RSV) for 48 h in the presence of RSV. The activity of the cells was measured by the CCK8 method (*n* = 3 biological replicates). **D**, RSV pharmacokinetic determination flowchart was divided into four groups, including Group 1 (RSV, 100 mg/kg, n = 8), Group 2 (*E. coli* BL21 (pET28a) 100 mg/kg, n = 9), Group 3 (*E. lenta* + RSV, 100 mg/kg, n = 8), and Group 4 (*E. coli* BL21 (RER) + RSV, 100 mg/kg, n = 8). **E**, RSV content in the serum of mice treated in different groups. **F**, Areas under the plasma RSV-concentration curves (AUC) in E. Data are presented as mean ± s.d. Error bars show s.d. Statistical analysis was performed by a two-tailed Student’s *t-*test. (n.s., not significant; *, p<0.05; **, p<0.01; ***, p<0.001; ****, p<0.0001).

### DHR demonstrates a mitigating effect on colitis in mice

Given that *E. lenta* J01 can metabolize RSV into DHR through RER catalysis to achieve higher biological activity, we further investigated the effect of *E. lenta* J01 and BL21 (RER) in animals. The mice used for the experiment were divided into four groups as follows: (1) the control group (RSV group), which received intragastric administration of phosphate-buffered saline (PBS) (100 μL/mouse) and RSV (100 mg/kg); (2) the *E. lenta* J01 group (*E. lenta* + RSV group), which received intragastric administration of *E. lenta* J01 (1×10^8^ CFU/100 μL/each) and RSV (100 mg/kg); (3) the BL21 (pET28a) group (BL21 + RSV group), which received intragastric administration of BL21 (pET28a) (1×10^8^ CFU/100 μL/each) and RSV (100 mg/kg); (4) the BL21 (RER) group (BL21 (RER) + RSV group), which received intragastric administration of BL21 (RER) (1×10^8^ CFU/100 μL/each) and RSV (100 mg/kg) (Fig. 4D). The pharmacokinetic analysis of RSV in the blood was then performed to evaluate the in vivo effect of *E. lenta* J01 and BL21 (RER). As depicted in Fig. 4E, the concentration of RSV reached the highest level at 30 min after intragastric administration and then gradually decreased in the RSV group and the BL21 + RSV group; in contrast, the RSV concentration decreased significantly at 30 min in the *E. lenta* + RSV group and the BL21 (RER) + RSV group. Consistent with these results, the areas under the RSV concentration curves (AUC) of the *E. lenta* + RSV group and the BL21 (RER) + RSV group were lower than those of the RSV group and the BL21 + RSV group (Fig. 4F), indicating that *E. lenta* J01 and BL21 (RER) had high RSV-metabolizing activity in vivo.

Next, we investigated the correlation between the metabolic effect of *E. lenta* J01 on RSV and inflammatory diseases, and chose a colitis mouse model for testing. To obtain a mouse model of acute colitis, dextran sulfate sodium (DSS) was used for induction treatment in mice. Then, the colitis mice were divided into six groups (Fig. 5A) as follows: (1) the DSS group, which received intragastric administration of 100 μL PBS; (2) the DSS + RSV group, which received intragastric administration of RSV (100 mg/kg); (3) the DSS + *E. lenta* group, which received intragastric administration of 100 μL *E. lenta* J01 (1×10^8^ CFU/mouse); (4) the DSS + *E. lenta* + RSV group, which received intragastric administration of RSV (100 mg/kg) and 100 μL *E. lenta* J01 (1×10^8^ CFU/mouse); (5) the DSS + BL21 + RSV group, which received intragastric administration of RSV (100 mg/kg) and 100 μL BL21 (pET28a) (1×10^8^ CFU/mouse); (6) the DSS + BL21(RER) + RSV group, which received intragastric administration of RSV (100 mg/kg) and BL21 (pET-28a-RER) (1×10^8^ CFU/mouse). The wild-type mice that received intragastric administration of PBS were set as the control group (Con) (Fig. 5A). We found that the RSV concentration in the internal contents (ileum, cecum, and colon) decreased more significantly in the DSS mice simultaneously exposed to RSV and *E. lenta* or BL21 (RER) (Figs. 5B[D). Concurrently, DHR formation was detected in the internal content of the colon in the abovementioned two groups (Fig. 5E), further validating the efficient conversion of RSV in these two groups.

**Fig. 5.**
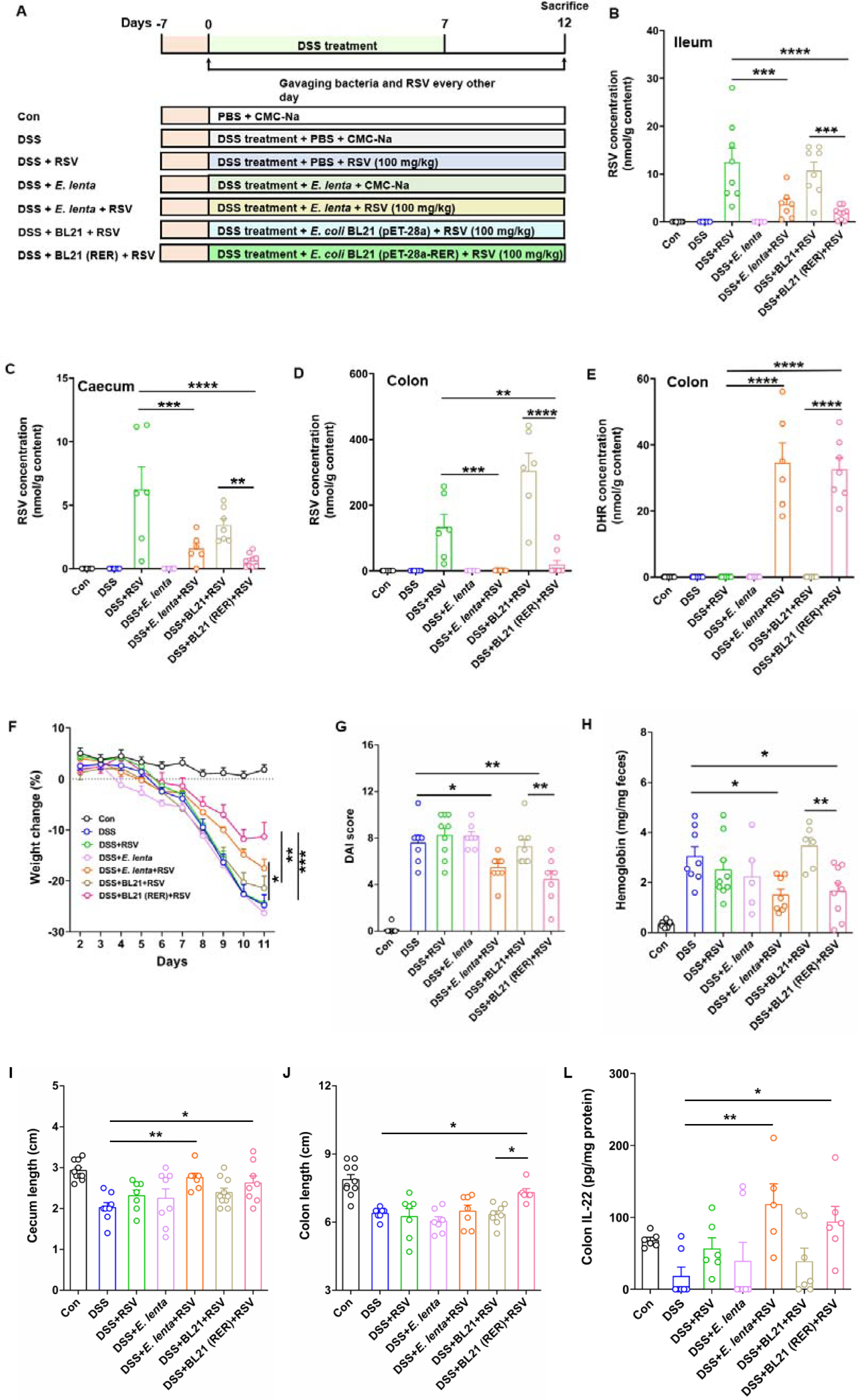
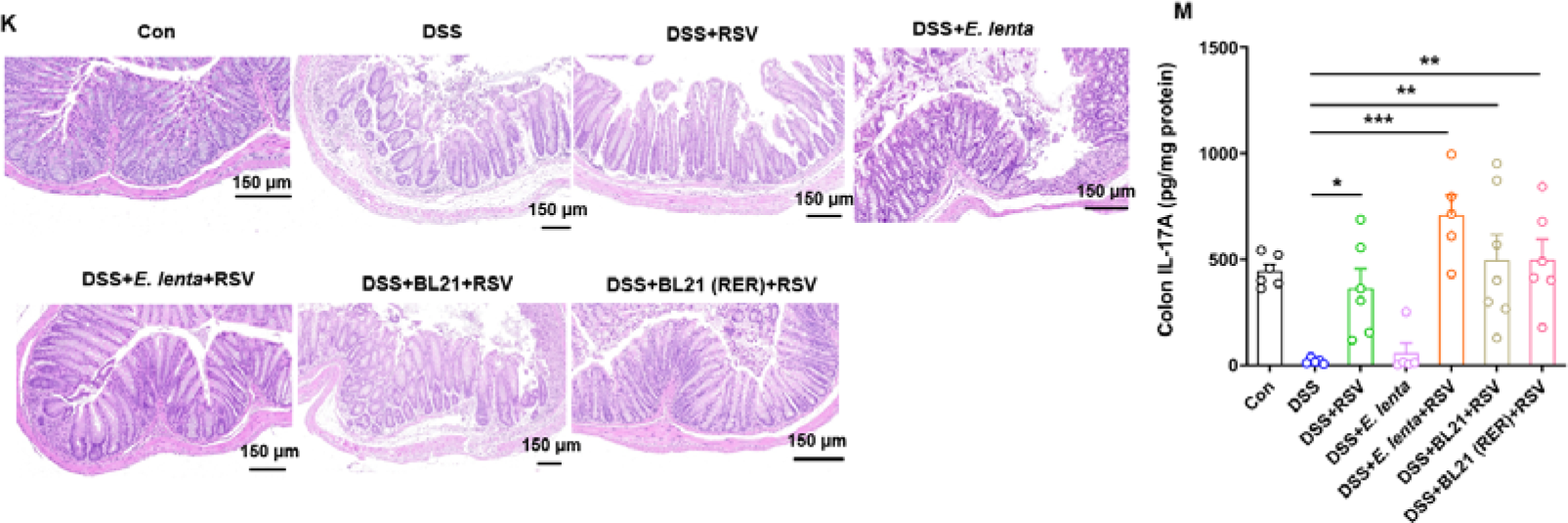
| Therapeutic effect of RSV, *E. lenta*, *E. lenta* + RSV, BL21 + RSV, and BL21 (RER) + RSV on DSS-induced colitis in mice. **A**, Schematic illustration of constructing a DSS-induced colitis mouse model and the oral supplementation of bacteria and RSV (Methods). The DSS-induced colitis mice were divided into six groups based on the different treatments, namely DSS, DSS + RSV, DSS + *E. lenta* J01, DSS + *E. lenta* J01 + RSV, DSS + BL21 + RSV, and DSS + BL21 (RER) + RSV. **B–D**, The RSV concentrations in the intestinal contents of ileum, cecum, and colon. **E**, The DHR concentration in the intestinal content of the colon. **F**, The body weight changes of different groups. **G**, The DAI scores of different groups (Day 9). **H**, The hemoglobin concentration in the feces of different groups (Day 7). **I–J**, The cecum and colon length of different groups. **K**, H&E staining of the colon tissue of different groups. **L–M**, The concentration of anti-inflammatory cytokines (IL-22 and IL-17A) in the colon of different groups. Data are presented as mean[±[s.d. (*n* ≥ 5). The statistical significance of data was determined using a one-way analysis of variance (*, p<0.05; **, p<0.01; ***, p<0.001; ****, p<0.0001).

Then, we evaluated the disease severity of colitis in different groups. As shown in Figs. 5F and 5G, the body weight change and disease activity index (DAI) score improved more significantly in the DSS + *E. lenta* + RSV group and the DSS + BL21 (RER) + RSV group than in the DSS group. Similarly, the concentration of hemoglobin in the fecal samples of the DSS + *E. lenta* + RSV group and the DSS + BL21 (RER) + RSV group decreased significantly (Fig. 5H). Furthermore, compared with the DSS group, the cecum was much longer in the DSS + *E. lenta* + RSV group and the DSS + BL21 (RER) + RSV group, while the colon was longer in the DSS + BL21 (RER) + RSV group (Figs. 5I and Figs. 5J). Histological examination of colon sections revealed more intact epithelial cells and regular crypt structure in the DSS + *E. lenta* + RSV group and the DSS + BL21 (RER) + RSV group (Fig. 5K). Given that the level of anti-inflammatory cytokines may be associated with the progression of colitis, we detected the level of anti-inflammatory cytokines IL-22 and IL-17A in the colon tissue of different groups. Compared with the DSS group, the concentration of IL-22 and IL-17A increased in the DSS + *E. lenta* + RSV group and the DSS + BL21 (RER) + RSV group (Figs. 5L–M). All these findings indicate that *E. lenta* and BL21 (RER) can metabolize RSV into DHR, thereby alleviating the disease severity of ulcerative colitis.

### DHR restores the disordered gut microbiota in the colitis mouse model

To further assess the effect of DHR on gut microbiota of colitis mice, we performed the 16S rRNA amplicon sequencing toward the different mice groups. The Shannon index did not significantly differ between the seven groups (Fig. 6A). As for the β-diversity, the gut microbiome of the colitis mice was quite different from that of the wild-type control mice (Fig. 6B). Notably, the gut microbiome of the mice from the DSS + *E. lenta* group and the DSS + *E. lenta* + RSV group differed from that of the other colitis groups. Analysis of the gut microorganisms in different groups revealed that the abundance of the colitis-causing genus *Escherichia-Shigella* decreased in the DSS + *E. lenta* + RSV group, while the abundance of the beneficial genera *Akkermansia* and *Faecalibaculum* increased in the DSS + *E. lenta* + RSV group and the DSS + BL21 (RER) + RSV group (Fig. 6C). Based on this finding, we performed correlation analysis to decipher the relationship of the relative abundance of gut microorganisms with the DAI score, hemoglobin concentration, and concentrations of anti-inflammatory factors IL-17A and IL-22. As shown in Fig. 6D, the concentrations of IL-17A and IL-22 positively correlated with the relative abundance of *Akkermansia*, while the relative abundance of *Faecalibaculum* showed a negative correlation with the DAI score and hemoglobin concentration. For the genus *Escherichia-Shigella*, the relative abundance negatively correlated with the concentrations of IL-17A and IL-22, and positively correlated with the DAI score and hemoglobin concentration (Fig. 6D). We also analyzed the correlation between the relative abundance of the *rer* gene and the concentrations of IL-17A and IL-22, and found that they positively correlated with each other (Figs. 6E-6F). These results indicate that DHR derived from RSV can benefit the colitis mice by restoring the disordered gut microbiota.

**Fig. 6.**
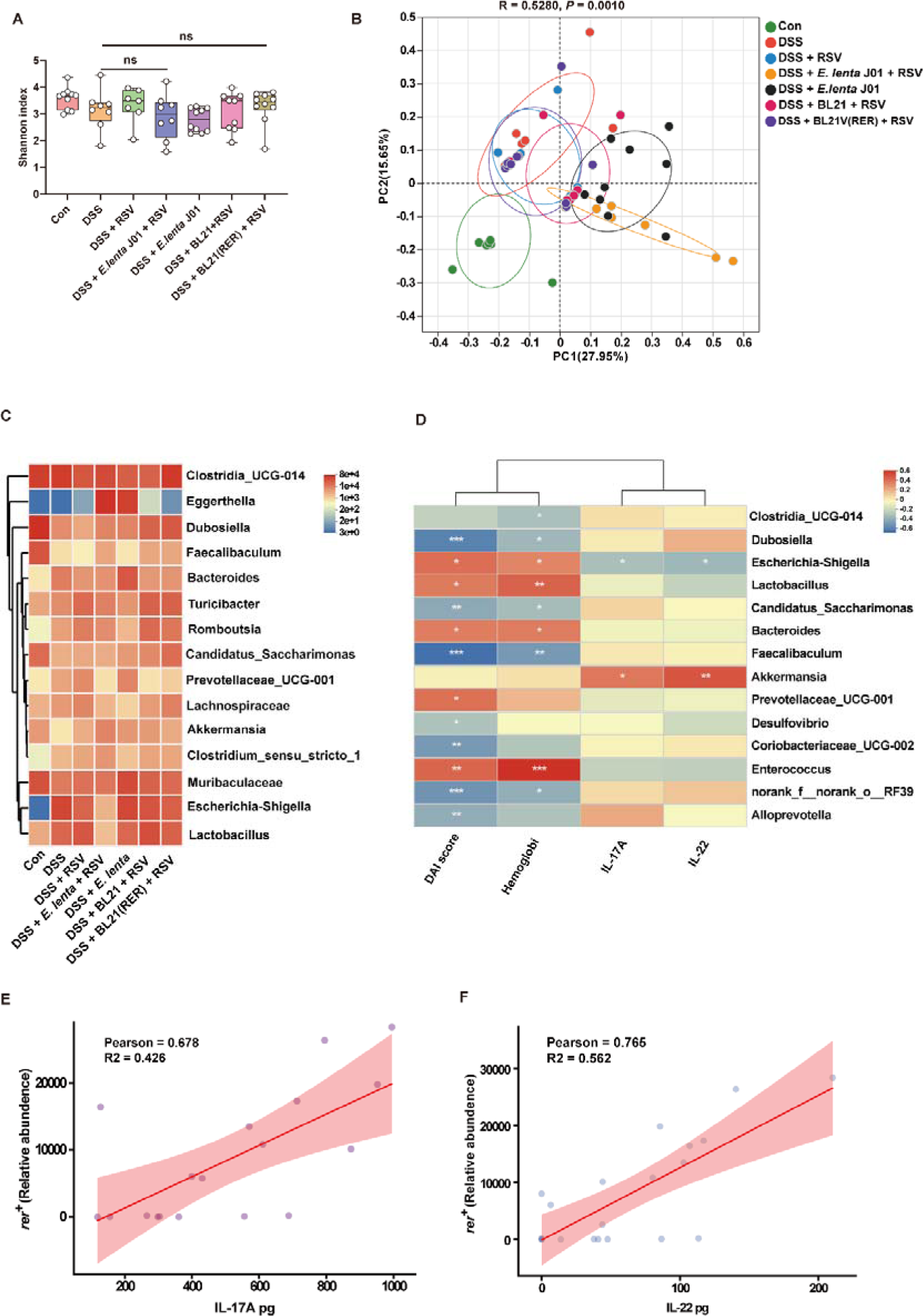
| RER regulates and restores the disordered gut microbiota of colitis mice by converting RSV to DHR. **A**, Assessment of α-diversity indexes. The genus-based Shannon indexes of the seven mice groups were assessed using faith effective counts. **B**, PCoA analysis based on the abundance of gut microbiota in seven groups of mice. ANOSIM (analysis of similarities), R = 0.5280, p = 0.001. **C**, Heatmap analysis of the top 15 distinguishing bacterial taxa between the seven mice groups. The horizontal coordinate represents different bacteria the vertical coordinate represents different groups, and the colors of the color blocks represent different relative abundance of bacteria. **D**, Correlation analysis of the relationship between the 14 bacterial genera and four colitis-related parameters. The differential bacteria in terms of relative abundance between seven groups were used to perform Pearson correlation with cytokines. **E–F**, We performed correlation analysis using the relative abundance of rer genes and cytokines (IL-17A and IL-22 concentrations). Data are presented as mean ± s.d. Error bar represents s.d. Statistical significance was determined by one-way ANOVA. (n.s., not significant; *, p<0.05; **, p<0.01; ***, p<0.001;****, p<0.0001).

### Distribution and abundance of RER in the gut microbiota of patients with colitis and healthy individuals

To investigate whether RER-like enzymes are widely distributed in the human gut microbiota, we analyzed 1,325 samples from healthy individuals and 334 samples from patients with colitis obtained from the GMrepo database, covering different countries, genders, and weights^16^. We found that *rer*^+^ bacteria were detected in 46.41% and 59.85% of patients with colitis and healthy individuals, respectively (Fig. 7A), indicating that the *rer* genes are widespread in intestinal microbiota. Of note, the relative abundance of *rer*^+^ bacteria in healthy individuals was much higher than that in patients with colitis (Fig. 7B). Analysis of different age and gender groups further revealed that the relative abundance of *rer*^+^ bacteria reached a maximum of 10% in the age group of 20[40 years old (Fig. 7C). In addition, the abundance of *rer*^+^ bacteria in the female intestine was significantly higher than that in male intestine (Fig. 7D, Fig. S9). Overall, our findings indicate that *rer*^+^ bacteria are widely distributed in human intestines, with an effect on converting RSV into the smaller molecule DHR with stronger biological activity.

**Fig. 7.**
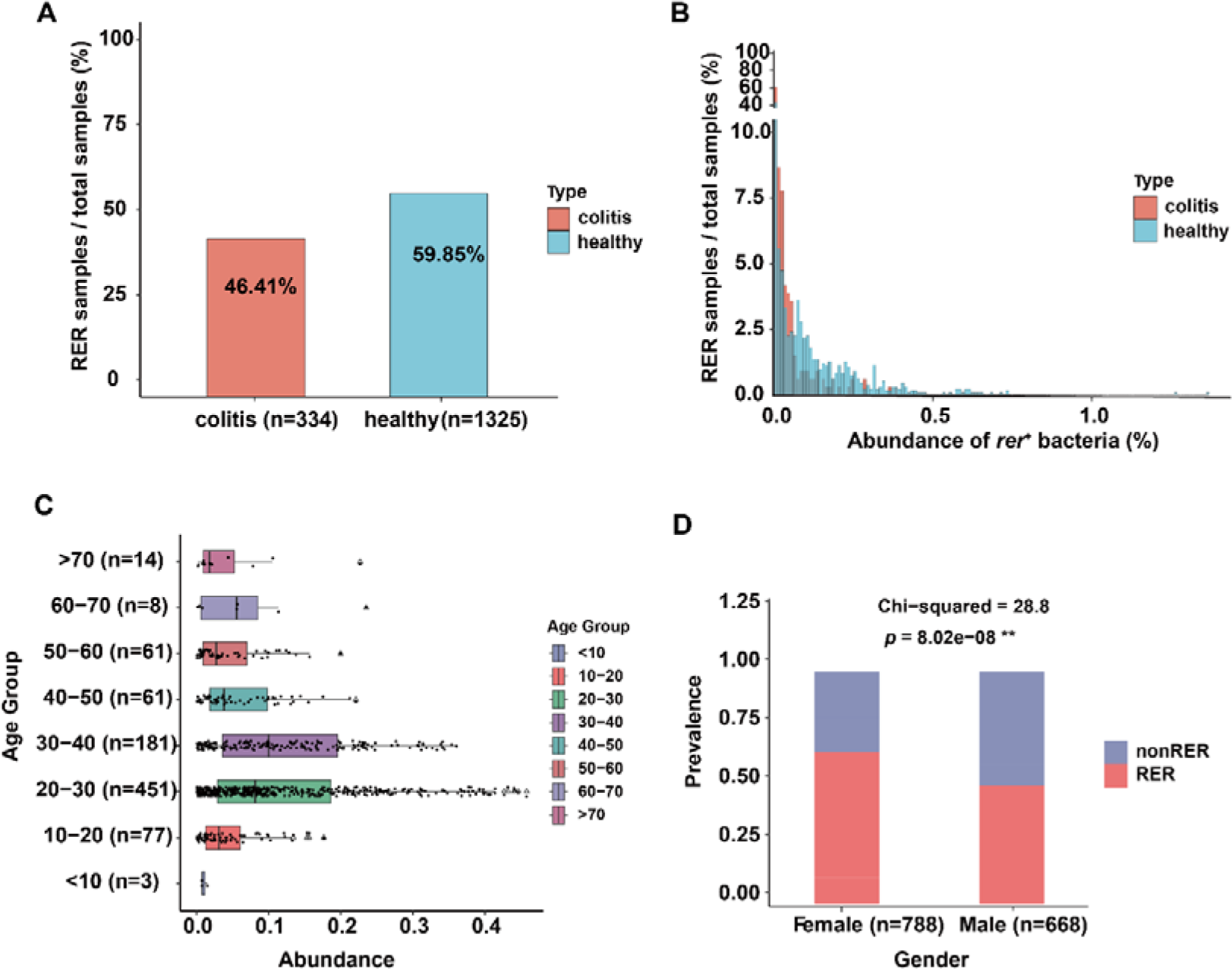
| Prevalence and abundance analysis of *rer*^+^ bacteria. **A, B**, Prevalence, and abundance of *rer*^+^ bacteria in the gut microbiota of healthy humans (blue, GMrepo, *n* = 1,325) and patients with colitis (red, this study, *n* = 334). **C**, The prevalence of *rer*^+^ bacteria among different age groups. The triangular symbols represent outliers. Values greater than the upper quartile plus 1.5 times the interquartile range, or less than the lower quartile minus 1.5 times the interquartile range were defined as outliers. **D**, The prevalence of *rer*^+^ bacteria in the male and female samples. The chi-squared test was performed between the two gender groups.

## Discussion

RSV is a highly active and efficacious plant-derived natural product. It has been attracting widespread attention given that it plays a pivotal role in the prevention and treatment of various diseases^17^. It has been shown that RSV can be metabolized in the human body, and its efficacy is closely linked to gut microbiota^10,18,19^. However, it remains unknown which gut microorganisms can efficiently metabolize RSV, and it is still unclear how the metabolic products affect the physiological state of the host. In the present study, we discovered that the intestinal bacterium *E. lenta* J01 encodes a kind of unreported RSV reductase, RER, which can efficiently convert RSV into DHR, thereby showcasing enhanced efficacy in inhibiting tumor-cell growth and improving intestinal inflammation. In addition, we found a significantly higher abundance of the *rer*-like genes in the intestinal bacteria of healthy individuals relative to the patients with enteritis. These findings expand our knowledge of the importance of intestinal microorganisms and ene-reductases to humans.

Thus far, very few intestinal bacteria (i.e., *S. equolifaciens* and *A. equolifaciens*) have been reported to metabolize RSV, and their metabolic efficiency is low (≤0.21 μM/h)^12^. In contrast, the *E. lenta* J01 identified in this study achieved an RSV-conversion rate of 20 μM/h—nearly 100 times the abovementioned level— demonstrating the existence of intestinal microorganisms capable of efficiently consuming RSV. As a major group of intestinal bacteria, *E. lenta* possesses unique metabolic characteristics and diverse chemical transformation capabilities, and it is capable of metabolizing various drugs (e.g., and levodopa)^3,20^ and dietary substrates (e.g., bile acids and plant lignans)^2,21^. Recent studies have revealed that the genome of *E. lenta* encodes numerous respiratory-like reductases, which facilitate energy production in this bacterium through electron receptors in the gut^22^. Since RSV has an unsaturated double bond, it can serve as an electron acceptor to support energy metabolism in *E. lenta*.

DHR is considered the main reductive product of RSV. It can be further converted into lunularin and 3,4’-dihydroxy-trans-stilbene through a dehydroxylation reaction in the human gut^12^. However, we did not detect the formation of these two chemicals by *E. lenta* J01 in the presence of RSV, suggesting that the conversion of RSV into these two products depends on other intestinal microorganisms. Therefore, it seems that the complete metabolism of RSV in the human gut requires the collaboration of multiple intestinal microorganisms.

Our animal experiments revealed that gavaging mice solely with RSV had no significant effect on the phenotype of colitis; however, simultaneous gavaging with *E. lenta*/BL21 (RER) and RSV led to the formation of DHR and an increase in the levels of the anti-inflammatory factors IL-22 and IL-17A in the mouse colon, thereby alleviating the phenotype of colitis (Figs. 5L–M). This suggests that in DSS-induced colitis mice, there was no microorganism capable of efficiently metabolizing RSV, or the abundance of such microorganisms in the gut was very low. Previous studies have shown that DHR exerts anti-colitis effects by activating the immune pathway aryl hydrocarbon receptor (AHR)^13^ and inhibiting signaling pathways related to intestinal cell autophagy and inflammation induction, such as SIRT1/mTOR^23^ and Wnt/β-catenin^24^. Considering that *E. lenta* J01 and BL21 (RER) efficiently metabolized RSV to generate DHR (Figs. 5B–E), we speculate that the anti-colitis effect produced by simultaneous gavaging with *E. lenta*/BL21 (RER) and RSV in this study may also be achieved through the abovementioned mechanism, which requires further validation.

Gavaging with higher doses of *Lactobacillus* can alleviate the phenotype of DSS-induced colitis by exerting an anti-inflammatory effect^25–27^. However, in the present study, there was a significant increase in the relative abundance of gut *Lactobacillus* in the DSS-treated group, which positively correlated with the DAI score and fecal hemoglobin content (an important indicator of fecal occult blood) in colitis mice (Figs. 6C–D), suggesting that some intestinal *Lactobacillus* strains may exacerbate colitis. This indicates that, when using *Lactobacillus* strains for colitis treatment, the metabolic characteristics and anti-inflammatory properties of specific strains should be carefully considered to avoid potential counterproductive effects.

In summary, we identified a new intestinal *E. lenta* strain and a novel class of ene-reductases, RER, with exceptional RSV-metabolizing activity. Both in vitro culturing and in vivo animal experiments suggest the contribution of *E. lenta*–mediated RSV transformation to its efficacy, thereby providing new insights into the interaction between RSV and intestinal microbiota. The RER-like enzymes may also serve as a potential catalyst for the biological production of DHR, an RSV derivative with stronger bioactivity. Additionally, considering that the complete degradation and conversion of RSV in the human body may require multiple intestinal microorganisms of coexistence, more comprehensive and in-depth research is required to elucidate the underlying mechanisms.

## Materials and Methods

### Fecal sample collection and pretreatment

Fresh fecal samples of healthy volunteers were collected and stored at [80 °C (≤ 30 min from collection to cryopreservation). Next, the frozen fecal samples (three grams) were brought into an anaerobic chamber (Whitley A35 Workstation, Don Whitley Scientific Limited, UK) and suspended in 25 mL of sterile phosphate buffer (8 g/L of NaCl, 0.2 g/L of KCl, 1.44 g/L of Na_2_HPO_4_, and 0.24 g/L of KH_2_HPO_4_) supplemented with 0.1% L-cysteine (PBSc) in a 50 mL sterile test tube. The insoluble particles were removed from the suspension by passing through a cell filter (SanGon Biotech, Shanghai, China). The filtrate was then mixed with an equal volume of glycerol (40%, vol/vol) dissolved in PBSc, placed in sealed glass vials, and frozen at [80 °C for the following experiments.

### Evaluation of RSV metabolism by the fecal microbiome

The *ex vivo* culturing of the gut microbiome from stool samples was carried out as described previously^28^. Briefly, 200 µL of each glycerol stock from the abovementioned fecal samples was inoculated into a slightly modified GAM Broth Medium (mGAM) (5 mL) for inoculum preparation. After 12 h of cultivation at 37 °C, 200 µL of culture was transferred into the medium (5 mL) supplemented with RSV (200 μM). The cultures were incubated at 37 °C for 24 h and then centrifuged at 14, 000 *g* for 10 min at 4 °C. The supernatant was used for HPLC analysis to determine the concentration of the residual RSV and its derivatives.

### Quantification of RSV and its derivative DHR

The concentrations of RSV and DHR were determined using high-performance liquid chromatography (HPLC) (Agilent 1260 infinity HPLC system) with a ZORBAX SB-C18 column (particle size, 5 μm; 250 mm × 4.6 mm) (Agilent Technologies, Santa Clara, USA) and a UV diode array detector (Agilent Technologies, Santa Clara, USA). All the compounds were confirmed by comparing their retention times and UV spectra to those of the standard substances. Solution A (methanol and 0.1% formate) and solution B (0.1% formate in water) served as the mobile phases to form a gradient as follows: 5% solution A for 2 min, 5–100% solution A for 12 min, and 100% solution A for 4 min. The flow rate was 1 mL[min^−1^. The concentrations of samples were calculated based on the peak area-based calibration curves of various concentrations of standards.

### Enrichment cultivation and isolation of the RSV-consuming strains

Human stool samples were pretreated as described above. The obtained supernatant (200 µL) was transferred into the mGAM^28^ medium containing 100 μM RSV. After incubating at 37°C for 12 h, passage to fresh mGAM medium for cultivation with a 5% inoculation volume. The culture of every successive passage was incubated for 24 h. Next, the culture was transferred into the mGAM medium containing different concentrations of RSV (80, 120, 160, 240, and 300 μM) for six rounds of subculturing. The final enrichment cultures were diluted to different concentrations and then streaked onto the BHI agar plates. After 24 h of incubation at 37 °C, colonies that appeared on the agar were picked and inoculated into the liquid mGAM medium containing 200 μM RSV. After 24 h of incubation at 37 °C, the culture was centrifuged at 14, 000 *g* for 10 min at 4 °C. The obtained supernatant was used for HPLC analysis to determine RSV concentration.

Once RSV in the medium was found to be consumed, the corresponding colony that displayed this activity was picked out for PCR amplification of the 16S rRNA gene using the primers 27F and 1492R (Table S7), yielding the target DNA fragment for sequencing. The gene sequence was analyzed using BLASTn (https://blast.ncbi.nlm.nih.gov/Blast.cgi). The identified *E. lenta* strain was then inoculated into the liquid mGAM medium (containing 200 μM RSV) to ensure its RSV-consuming ability. Simultaneously, the grown *E. lenta* cells were harvested by centrifugation at 4, 000 *g* for 10 min at 4 °C and then stored at [20 °C for the following experiments.

### Cancer cell culture assays

The HCT116 human colorectal adenocarcinoma (CRC) cell line was procured from the American Type Culture Collection (ATCC). The cells were cultured in Dulbecco’s modified Eagle’s medium (DMEM) (Gibco BRL.Grand Island, NY, USA), supplemented with 10% fetal bovine serum (FBS, YOUSHI, Wuhan, China), and incubated at 37°C in a humidified incubator with 5% CO_2_. Cell passage was conducted using trypsin–EDTA (Gibco, Burlington, Canada). To assess the inhibitory activity of RSV and its metabolites on HT116 cells, initially 200 μM RSV was introduced into the main medium for the control group, while the treatment group was additionally exposed to the *E. lenta* J01 strain. Subsequently, both groups were incubated in an anaerobic workstation for 24 h. Next, the culture medium samples were collected, subjected to centrifugation at 12,000 rpm to eliminate bacteria, and subsequently sterilized through a 0.22-μm filter membrane, followed by the detection of RSV and DHR using HPLC. Following this, the treated RSV and its metabolites were introduced into the HT116 cell culture plate. Cellular activity of HCT116 was assessed using the CCK8 method after 72 h of cultivation^29^.

### Construction of DSS-induced murine model

Male C57BL/6J mice (6–8 weeks) were purchased from Charles River Laboratories China. The animals were maintained under specific pathogen-free conditions on a 12 light–dark cycle at a temperature of 20[°C and 40–60% humidity, with free access to food and water. All experiments were performed according to the ethical guidelines of the CAS Center for Excellence in Brain Science and Intelligence Technology, Chinese Academy of Sciences (ethical review approval number NA-041-2022). After a week of acclimatization period, the mice were randomly divided into seven groups. The construction of the DSS-induced murine IBD model was performed by feeding the mice with drinking water containing 3.5% DSS (MB5535, Meilun Bio) for 7 days. The mice in the control group (Con group) received normal water. Then, the DSS-induced colitis mice were divided into six groups and gavaged with PBS, RSV, *E. lenta*, *E. lenta* + RSV, BL21 + RSV, BL21 (RER) + RSV once per day for twelve days (1×10^8^ CFU/100 μL/mice). The mice in the control group (Con group) were gavaged with PBS. During the experiment, the weight of the mice was recorded every day. The DAI score^30^ and fecal hemoglobin concentration were measured on day 9 and day 7, separately. On the 12th day, the mice were sacrificed to collect the distal portions of colon tissues. Then, the colon tissues were used to perform the hematoxylin and eosin (H&E) staining and anti-inflammatory cytokines (IL-22 and IL-17A) detection.

### Pharmacokinetics experiments

Male C57BL/6J mice (6–8 weeks) were intragastrically administered with RSV (100 mg/kg) at the same time. Then, the mice were divided into four groups and gavaged with RSV (RSV group); *E. coli* BL21 (pET28a) + RSV (BL21 + RSV group); *E. lenta* J01+ RSV (*E. lenta* + RSV group), and *E. coli* BL21 (pRER) + RSV (BL21 (RER) + RSV group) one h later (CFU=1×10^8^/mice). The resveratrol concentration in the whole blood was assayed with HPLC at 0, 1, 2, 3, and 4 h post-gavage.

### Histology assay

For H&E staining, the distal colons of mice were collected and fixed in 4% paraformaldehyde for 24 h. Then, the colon tissue was embedded in paraffin, and sectioned for H&E staining. Finally, the mounted sections were imaged using a Pannoramic MIDI slide scanner (3D HISTECH).

### Colon tissue cytokine detection

To detect the concentration of anti-inflammatory cytokines IL-22 and IL-17A, the colon tissues (100 mg) were weighed and dissolved in 500 μl PBS buffer. After homogenization treatment, the samples were centrifuged at 12,000 rpm for 15 min (4°C). Then, the supernatants were collected and used to measure the concentration of total protein and anti-inflammatory cytokines. The protein concentration was assayed according to the manufacturer’s instructions for the Omni-Easy™ Ready-to-Use BCA Protein Quantification Kit (Yamei, ZJ102L). The concentration of IL-17A and IL-22 in colon tissue was assayed by using the mouse IL-17A (Proteinintech, KE10020) and IL-22 (Proteinintech, KE10041) ELISA kits according to the instructions.

### Fecal hemoglobin detection

The feces (100mg) were weighed and dissolved in 100 μl PBS buffer. After homogenization treatment, the samples were centrifuged at 12,000 rpm for 15 min (4°C). Then, the hemoglobin concentration in the supernatants was assayed according to the manufacturer’s instructions for the hemoglobin colorimetric assay kit (Beyotime, P0381S).

### RNA sequencing and data analysis

*E. lenta* strains were grown in mGAM medium with RSV (200 μM RSV, *n* = 2) or without RSV (equal proportions of methanol, *n* = 2) when they reached the logarithmic metaphase. After coincubation for 2 h, the cells were centrifuged at 4,000 *g* and 4°C for 10 min and frozen immediately with liquid nitrogen.

Total RNA was extracted from the tissue using TRIzol® Reagent in accordance with the manufacturer’s instructions (Invitrogen), and genomic DNA was removed using DNase I (Takara). Then, RNA quality was determined using 2100 Bioanalyser (Agilent) and quantified using the ND-2000 (NanoDrop Technologies). High-quality RNA sample (OD_260/280_=1.8–2.2, OD_260/230_ ≥ 2.0, RIN ≥ 6.5, 28S:18S ≥ 1.0, >10 μg) was used to construct sequencing library.

RNA-sequencing strand-specific libraries were prepared following the TruSeq RNA sample preparation kit from Illumina (San Diego, CA), using 5 μg of total RNA. Shortly, rRNA removed by RiboZero rRNA removal kit (Epicenter) was fragmented using fragmentation buffer. cDNA synthesis, end repair, A-base addition, and ligation of the Illumina-indexed adaptors were performed in line with the Illumina’s protocol. Libraries were then size-selected for cDNA target fragments of 200–300 bp on 2% Low Range Ultra Agarose followed by PCR amplification using Phusion DNA polymerase (NEB) for 15 PCR cycles. After quantification by TBS380, paired-end libraries were sequenced by Illumina NovaSeq 6000 sequencing (Shanghai BIOZERON Co., Ltd.). The raw paired-end reads underwent trimming and quality control using Trimmomatic (SLIDINGWINDOW:4:15 MINLEN:75) (version 0.36 http://www.usadellab.org/cms/uploads/supplementary/Trimmomatic). Then, clean reads were separately aligned to the reference genome with orientation mode using the Rockhopper software (http://cs.wellesley.edu/~btjaden/Rockhopper/).

To identify differentially expressed genes (DEGs) between two samples, transcript expression levels were computed by the fragments per kilobase of read per million mapped reads (RPKM) method. Differential expression analysis was conducted using edgeR (https://bioconductor.org/packages/release/bioc/html/edgeR.html). The DEGs between the two samples were selected based on the following criteria: the logarithm fold-change ≥2 and the false-discovery rate (FDR) ≤0.05. To understand the functions of the DEGs, GO functional enrichment and KEGG pathway analyses were performed using GOATOOLS (https://github.com/tanghaibao/Goatools) and KOBAS (http://kobas.cbi.pku.edu.cn/kobas3), respectively. The DEGs were significantly enriched in the GO terms and metabolic pathways when their Bonferroni-corrected P value was lower than 0.05.

### Construction of plasmids

All the primers, plasmids, and strains used in this study are listed in Table S7.

The vector expressing the gene (LUA64_RS01375) coding for the RER protein from *E. lenta* was constructed as follows. The DNA fragment of the gene was obtained by PCR amplification using the *E. lenta* genomic DNA as the template and the primers LUA64_RS01375-f/ LUA64_RS01375-r. The PCR product was analyzed by agarose gel electrophoresis and the target band was recovered using the DNA gel recovery kit (Axygen Biotechnology Company Limited, Hangzhou, China). The yielded DNA fragment was linked with the linear plasmid pET28a vector (digested with *Nde*I and *Xho*I) using a Hieff Clone Plus One Cloning Kit (10912ES10, Yeasen, China), yielding the target plasmid pET28a-LUA64_RS01375.

The plasmids expressing the other genes (Table S8) from *E. lenta* were constructed with the same steps except for the primers used in PCR amplification.

### Protein production and purification

The plasmids harboring the coding sequences of target proteins were transformed into *E. coli* BL21 strain (DE3) for protein production. Transformants were grown on agar plates (LB medium containing 50 μg/mL of kanamycin) at 37 °C. Colonies on agar plates were then inoculated into liquid LB media (containing 50 μg/mL kanamycin) for cultivation at 37 °C for 12 h. The grown cells (2 mL) were inoculated into 200 mL of liquid LB medium (containing 50 μg/mL of kanamycin) for further cultivation at 37 °C. When biomass (OD_600_) reached ∼0.8, the protein expression was induced at 16 °C with the supplementation of 1 mM IPTG. After 16 h of incubation, *E. coli* cells were harvested by centrifugation at 5, 000 *g* for 10 min at 4 °C, and then resuspended in buffer (50 mM Tris-HCl, 100 mM NaCl, 10% glycerol, pH 7.9). Cells were lysed using a cell disruptor (French Press, Constant Systems Limited, UK), and the lysate was clarified by centrifugation at 12, 000 *g* for 1 h at 4 °C. The supernatant was loaded onto a Ni^2+^ Sepharose™ 6 fast-flow agarose column (GE Healthcare, Waukesha, WI, USA) for purification. The column was then washed with 150 mL protein solution A (20 mM Tris-HCl, pH 7.9, 10% glycerol, 500 mM KCl, 10 mM imidazole) followed by 150 mL protein solution B (20 mM Tris-HCl, pH 7.9, 10% glycerol, 500 mM KCl, 25 mM imidazole) and 300 mL protein solution C (20 mM Tris-HCl, pH 7.9, 10% glycerol, 500 mM KCl, 50 mM imidazole). The eluted fractions from solution C (containing target proteins) were applied to an Amicon Ultra 15 Centrifugal Filter (Millipore Billerica MA) for desalting and imidazole removal by using a buffer containing 100 mM Tris–HCl (pH 7.9), 200 mM NaCl, 10% (v/v) glycerol, and 3 mM DTT. The purified protein was stored in 50% (w/v) glycerol at ■80 °C.

### Liquid chromatograph mass spectrometry analysis to identify RSV and its derivatives

The RSV derivatives were obtained through the following three ways: (i) Coincubation of the *E. lenta* strain and RSV for 12 h. The RSV metabolite M was collected via HPLC. (ii) Coincubation of the RER enzyme and RSV for 30[60 min. The enzymatic product M was also collected using HPLC. (iii) Fecal samples from mice that had been administered RSV via gavage. The processing of fecal samples followed the methodology outlined in a prior report^15^.

The above samples were analyzed by Q Exactive quadrupole orbitrap high-resolution mass spectrometry coupled with a Dionex Ultimate 3000 RSLC (HPG) ultra-performance liquid chromatography (UPLC-Q-Orbitrap-HRMS) system (Thermo Fisher Scientific), with a HESI ionization source. The injection volume was 2 µL. Samples were separated with a Poroshell SB-Aq column (100 mm × 3.0 mm, 2.7 μm particle size; Waters). The mobile phase consisted of 0.1% formic acid in water (A), and acetonitrile (B). The gradient elution was set as follows: 0.0–8.0 min, 10%–95% B; 8.0–9.0 min, 95% B; 9.1–11.0 min, 10% B. The flow rate was 0.35 mL/min, and the column temperature was 40°C.

All MS experiments were performed in positive and negative ion modes using a heated ESI source. The applied source and ion transfer parameters were as follows: spray voltage of 3.5 kV (positive). In the ionization mode, the sheath gas, auxiliary gas, capillary temperature, and heater temperature were maintained at 40, 10 (arbitrary units), 300°C, and 350°C, respectively. The S-Lens RF level was set at 50. The Orbitrap mass analyzer was operated at a resolving power of 70,000 in full-scan mode (scan range: 100–1,200 m/z; automatic gain control (AGC) target: 1e6) and 175.00 in the Top 4 data-dependent MS2 mode (stepped normalized collision energy: 15, 30, and 45; injection time: 50 ms; isolation window: 1.5 m/z; AGC target: 1e5) with a dynamic exclusion setting of 4.0 s.

### RER enzyme assays and kinetics

The buffer for enzyme activity assays contained 100 mM Tris-HCl (pH 7.9), 100 mM NaCl, 10% (v/v) glycerol, FAD (0.03 mM at final concentration), and reduced methyl viologen (0.68 mM at final concentration). The reaction mixture (200 μL), containing 0.15 mg/mL RER, 100 μM of substrates (RSV, trans-stilbene, pterostilbene, and piceatannol, cis-resveratrol, naringenin, and p-hydroxycinnamic acid, apigenin, isoliquiritigenin, and chlorogenic acid), and buffer, was incubated at 37 °C for 30 min. The reaction was terminated by adding an equal proportion of methanol. The reaction mixture was then centrifuged at 14, 000 *g* for 10 min at 4 °C, and the supernatant was subject to HPLC analysis to determine the consumption of substrates. Each experiment was repeated in triplicate.

To determine the *K*_m_ and *k*_cat_ values of RER for RSV, the enzyme was tested at a concentration ([E]) of 0.15 mg/mL with various concentrations ([S]) of the substrates (RSV) in the reaction system. The reaction mixtures (200 μL) that contained 100 mM Tris-HCl (pH 7.9), 100 mM NaCl, 10% (vol/vol) glycerol, 0.03 mM FAD, 0.68 mM reduced methyl viologen, 0.15 mg/mL μM purified RER, and different concentrations of RSV (50, 100, 150, 200, 400, 600, 800, 1,000, 1,200, 1,400, 1,800, 2,000, and 3,000 µM) were incubated at 37 °C for 30 min. The reaction was terminated by adding an equal proportion of methanol. The reaction mixture was then centrifuged at 14, 000 *g* for 10 min at 4 °C. The supernatant was used to determine the RSV consumption by HPLC. The determination of the *K*_m_ and *k*_cat_ values of RER for DHR was carried out via the same steps, except a modified reaction mixture that contained 100 mM Tris-HCl (pH 7.9), 0.3 mM FAD, 0.15 mg/mL purified RER, and different concentrations of naringenin (100, 200, 400, 600, 800, 1,000, 1,200, 1,400, and 1,600 μM). Initial rates of substrate consumption (V) were obtained by linear regression for the data points within the initial 0–30 min reaction time (the substrate consumption versus time). The observed rate constant (*k*_obs_) was calculated by *k*_obs_ = V/[E] and then fit the equation (*k*_obs_= *k*_cat_* [S]/(*K*_m_ + [S]) in Graphpad Prism, yielding *K*_m_ and *k*_cat_ values.

### 16S rRNA sequencing and data analysis

The genomic DNA was extracted from microbial samples (a total of 16 culture samples and 62 samples of mice fecal contents) using the E.Z.N.A.® soil DNA Kit (Omega Bio-tek, Norcross, GA, U.S.) in accordance with the manufacturer’s instructions. The quality and concentration of DNA were determined by 1.0% agarose gel electrophoresis and a NanoDrop2000 spectrophotometer (Thermo Scientific, United States). The hypervariable region V3–V4 of the bacterial 16S rRNA gene was amplified using the primers 338F (5’-ACTCCTACGGGAGGCAGCAG-3’)/ 806R (5’-GGACTACHVGGGTWTCTAAT-3’) by using a T100 Thermal Cycler PCR thermocycler (BIO-RAD, USA). The PCR reaction mixture (20 µL) included 4 μL of 5 × Fast Pfu buffer, 2 μL of 2.5 mM dNTPs, 0.8 μL of each primer (5 μM), 0.4 μL of Fast Pfu polymerase, and 10 ng of template DNA. PCR amplification cycling conditions were as follows: initial denaturation at 95°C for 3 min, followed by 27 cycles of denaturing at 95°C for 30 s, annealing at 55°C for 30 s, and extension at 72°C for 45 s, and single extension at 72°C for 10 min, and end at 4°C. The PCR product was extracted from 2% agarose gel and purified using the PCR Clean-Up Kit (YuHua, Shanghai, China) in accordance with the manufacturer’s instructions, and quantified using Qubit 4.0 (Thermo Fisher Scientific, USA).

### Illumina PE300/PE250 sequencing

The purified amplicons were pooled in equimolar amounts and paired-end sequenced on an Illumina PE300 platform (Illumina, San Diego, USA) in accordance with the standard protocols by Majorbio Bio-Pharm Technology Co., Ltd. (Shanghai, China).

### Amplicon sequence processing and analysis

After demultiplexing, the resulting sequences were quality-filtered with fast (0.19.6) and merged with FLAS. Then, the high-quality sequences were de-noised using DADA2 plugin in the Qiime2 (version 2020.2) pipeline with recommended parameters, which obtains single-nucleotide resolution based on error profiles within samples. DADA2 denoised sequences are usually called amplicon sequence variants (ASVs). To minimize the effects of sequencing depth on alpha and beta diversity measures, the number of sequences from each sample was rarefied to 20,000, which still yielded an average Good’s coverage of 97.90%. Taxonomic assignment of ASVs was performed using the Naive Bayes consensus taxonomy classifier implemented in Qiime2 and the SILVA 16S rRNA database (v138).

Based on the ASVs information, rarefaction curves, and alpha diversity indexes including observed ASVs, the Shannon index was calculated with Mothur v1.30.1. The similarity among the microbial communities in different samples was determined by principal coordinate analysis (PCoA) based on Bray–Curtis dissimilarity using the Vegan v2.5-3 package. The PERMANOVA test was used to assess the percentage of variation explained by the treatment along with its statistical significance using the Vegan v2.5-3 package. The linear discriminant analysis (LDA) effect size (LEfSe) (http://huttenhower.sph.harvard.edu/LEfSe) was performed to identify the significantly abundant taxa (phylum to genera) of bacteria among the different groups (LDA score > 2, P < 0.05).

### Evaluation of the prevalence and abundance of microorganisms containing the RER-like proteins

To assess the prevalence and abundance of microorganisms carrying RER protein in the gut microbiota of both healthy individuals and patients with colitis, we identified microbial taxa containing RER protein. The RER protein sequence was employed as a query, and a homology search was conducted using Blastp against the nonredundant protein database in the NCBI. Microbial taxa with a protein sequence identity ≥40% were selected for further analysis. Subsequently, gut microbiota metagenomic data and metadata from both healthy individuals and patients with colitis were downloaded from the GMrepo v2 (ref. Dai, D. et al. Nucleic Acids Res (2022)) database. The criteria for sample selection encompassed the data type being metagenomic, the recent absence of antibiotic usage, and the successful completion of quality control. Specifically, for healthy samples, the criteria were disease = ‘D006262’ AND (experiment_type = ‘Metagenomics’ AND Recent. Antibiotics. Use = ‘N’ AND QC Status = 1), resulting in a total of 1,325 samples. For colitis samples, the criteria were disease = ‘D003093’ AND (experiment_type = ‘Metagenomics’ AND Recent. Antibiotics. Use = ‘N’ AND QC Status = 1), yielding a total of 334 samples. Subsequently, a custom script was employed to conduct a statistical analysis on the distribution of prevalence and abundance of microorganisms carrying RER proteins in the gut microbiota of both healthy individuals and patients with colitis.

### Construction of SSN

To establish an ene-reductases similarity network, a custom script was initially utilized to retrieve 6,473 amino acid sequences of predicted or characterized ene-reductases from the KEGG database. This dataset includes 1,379 sequences from the OYE family of oxido-reductases (OYE, EC 1.6.99.1), 221 sequences of oxygen-sensitive FAD- and [4Fe-4S]-containing enoate reductases (EnoR, EC 1.3.1.31), 3,114 sequences of medium-chain dehydrogenases/reductases (MDR, EC 1.3.1.74 and 1.3.1.48), 1,233 sequences of the short-chain dehydrogenase/reductase salutaridine/menthone reductase-like subfamily (SDR, EC 1.1.1.208), 526 sequences of NADPH-dependent quinone reductases (QnoR, EC:1.6.5.5). Homologous sequences of RER and FLR (Access ID: ANU40626.1) were chosen through alignment with the nonredundant protein sequence database at the National Center for Biotechnology Information (NCBI). A total of 63 RER sequences were chosen based on coverage exceeding 99% and identity values surpassing 56%. Similarly, 280 FLR sequences were selected based on coverage exceeding 80% and identity values surpassing 60%. Subsequently, the EFI-ENZYME SIMILARITY TOOL (https://efi.igb.illinois.edu/efi-est/) was used to construct a protein similarity network using the downloaded amino acid sequences. An alignment score cutoff of 10^−40^ resulted in an SSN with 4,089 nodes (with 95% identity) and 1,748,059 edges. Visualization and editing of the SSN were performed using Cytoscape 3.10.1, applying a layout through Prefuse Force Directed OpenCL Layout.

### Construction of the phylogenetic tree for *Eggerthella lenta* and the RER proteins

The genomic sequences of all of the *Eggerthella lenta* strains were retrieved and downloaded from the NCBI database. The dataset comprised both complete genomes and metagenome-assembled genomes (MAGs). The construction of the phylogenetic tree was conducted using the PhyloPhlAn tool (V3.0.67).

The identification of species-specific marker genes was executed using the built-in command “phylophlan_setup_database.py” in PhyloPhlAn 3.0. Default parameters were employed, utilizing 400 available universal marker genes to ensure the construction of phylogenies at a higher resolution. Specifically, the parameters used were “-diversity low --fast --min_num_marker 50,” signifying the exclusion of genomes with fewer than 50 identified marker genes. Other software main parameters applied in PhyloPhlAn 3.0 were as follows: (i) Diamond (v0.9.19.120): blastx –quiet --threads 1 --outfit 6 --more-sensitive --id 50 --max-hips 35 -k 0 --query-encode 11; (ii) Mafft (v7.505): --quiet --anysymbol --thread 1 –auto; (iii) FastTreeMP (v2.1.11): - quiet -pseudo -spr 4 -mlacc 2 -slownni -fastest -no2nd -mlnni 4 -lg; (iv) raxmlHPC-PTHREADS-SSE3 (v8.2.12): -p 1989 -m PROTCATLG. The resulting tree file underwent editing using iTOL (Interactive Tree of Life).

To construct the phylogenetic tree for RER proteins, a sequence alignment was initiated through a blastp comparison of the RER protein sequences against the NCBI nonredundant protein database. After that, we selected 63 RER protein sequences with a coverage greater than 59% and an identity value exceeding 56%. The homologous RER protein sequences identified were aligned using Mafft (v7.505). Following alignment, gaps and ambiguously aligned regions were removed through trimming. Subsequently, the phylogenetic tree was constructed using the FastTree program. The resulting tree file was then edited for visualization and analysis using iTOL.

## Data availability

Data supporting the findings of this study are available within the paper and the Supplementary Information files. *E. lenta* J01 genome sequencing raw data is available through NCBI-Genome associated with NCBI-Bioproject accession PRJNA1076688. All RNA-seq raw data is available through NCBI-SRA associated with NCBI-Bioproject accession PRJNA1077243. All 16S rRNA sequencing raw data is available through NCBI-SRA associated with NCBI-Bioproject accession PRJNA1076899 and PRJNA1076819. Datasets and strains generated and analyzed in the study are available from the corresponding author upon reasonable requests.

## Code availability

The amino acid sequences of enzymes (containing EC numbers) for SSN analysis were downloaded from KEGG. The generated source code is available at https://github.com/huiz0916/RER_gut

## Corresponding authors

Correspondence to Yunpeng Yang, Yang Gu, or Weihong Jiang.

## Acknowledgments

This work was supported by grants from the National Natural Science Foundation of China (no. 31921006), the 111 Project D18007 and a Project Funded by the Priority Academic Program Development of Jiangsu Higher Education Institutions (PAPD). We are grateful to Xiaoyan Xu and Yining Liu for HPLC and LC-MS/MS analysis.

## Author Contributions

W.J., Y.G., Y.Y., and Z.D. conceived and initiated the project; Z.D. and P.Y. performed the biochemical experiments; Y.Y., P.Y., and Q.S. performed the animal experiments; W.J., Y.G., Z.D., Y.Y., and P.Y. wrote the manuscript.

---

**Fig. S1.**
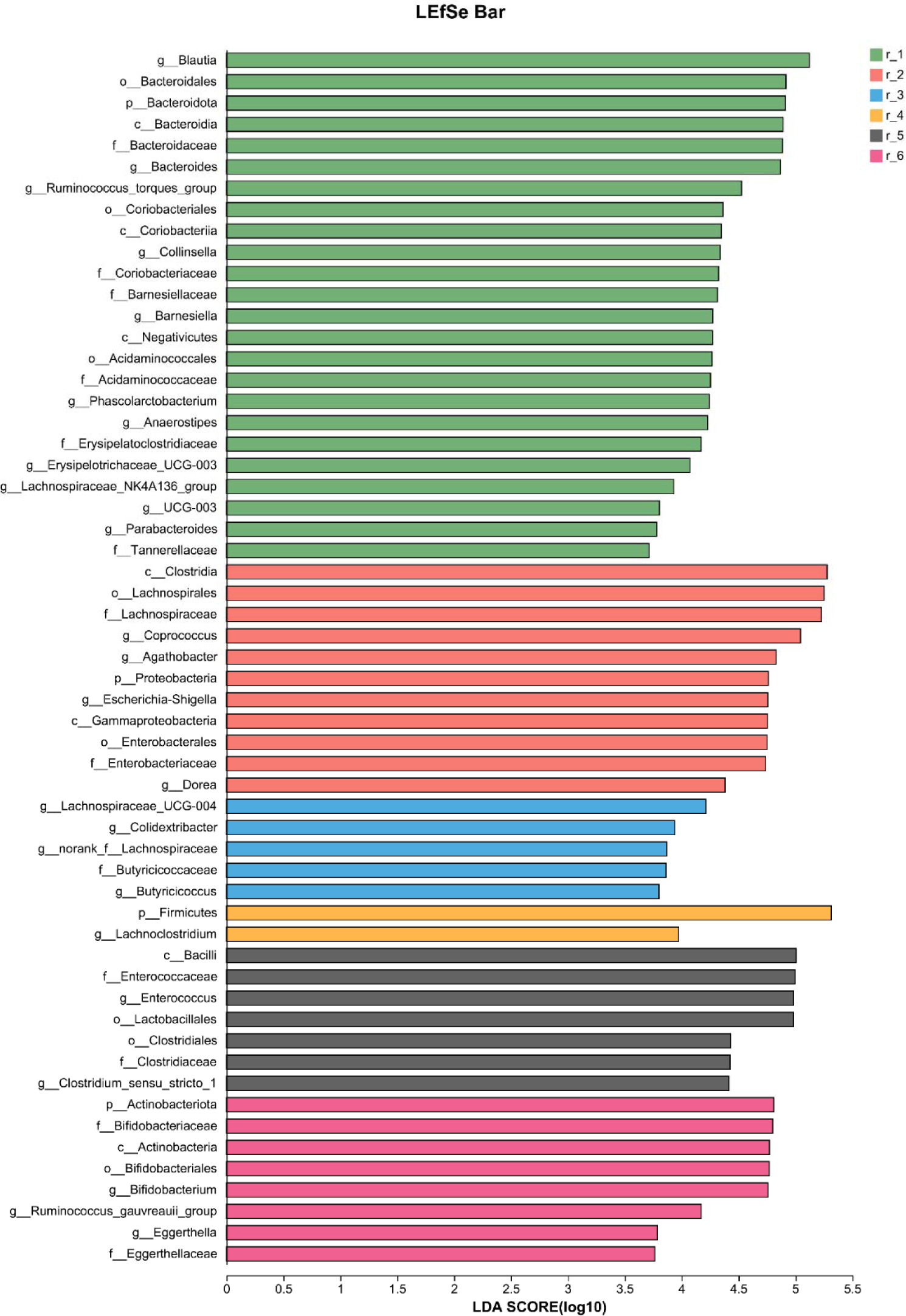
LEfSe analysis revealed the strains with potential RSV-metabolizing activity after enrichment culturing. A plot depicting LDA scores generated through the LEfSe method illustrates the differential abundance of specified taxa within the microbiomes of enrichment cultures associated with RSV enrichment.

**Fig. S2.**
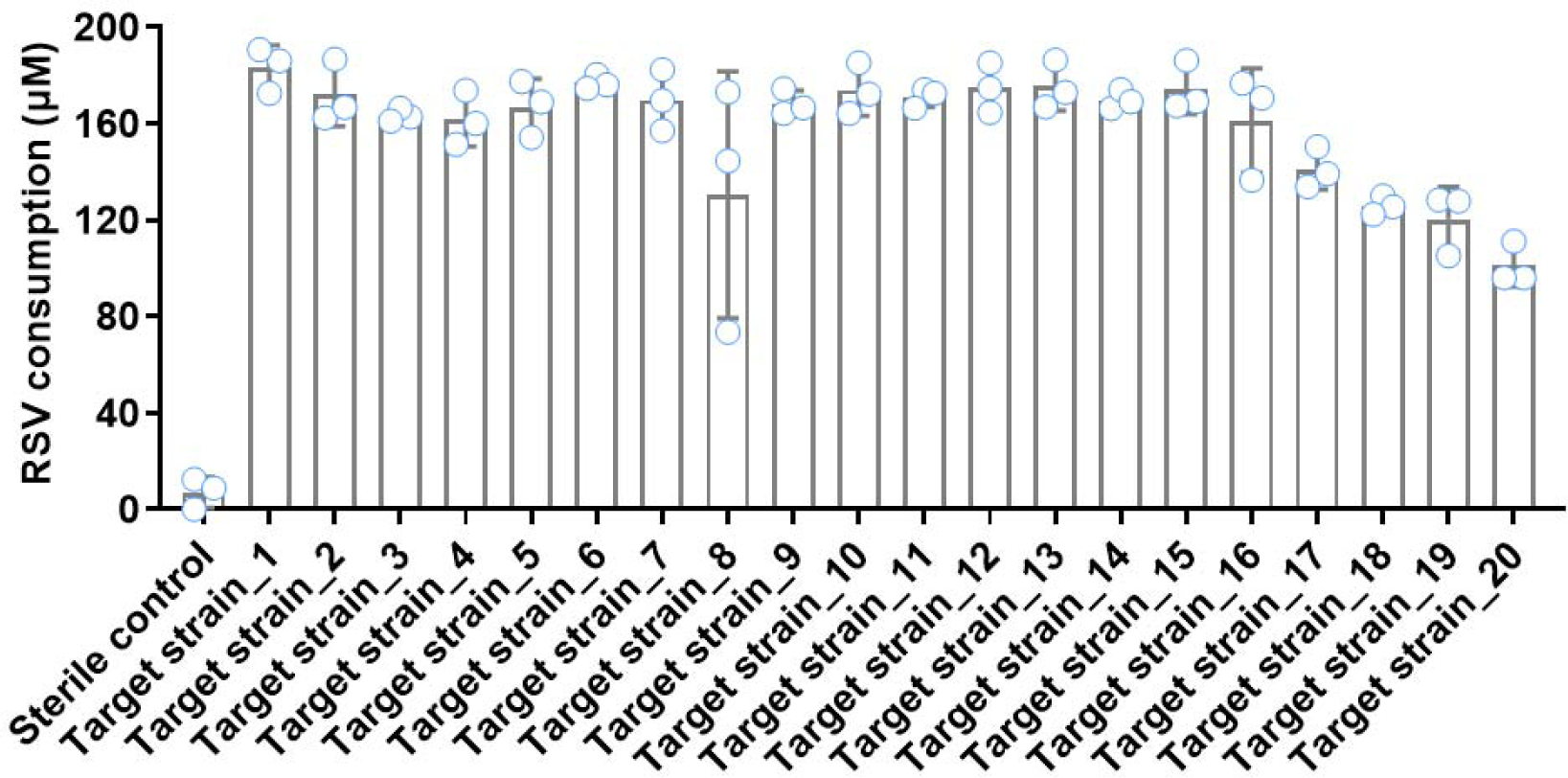
Identification of the RSV-metabolizing activity of the strains obtained from enrichment culturing. The enrichment culturing was performed anaerobically using the liquid mGAM medium. The residual RSV concentration in the medium (12 h cultivation) was determined by HPLC analysis. Data are represented as mean ± s.d. (*n* = 3). Error bars show s.d.

**Fig. S3.**
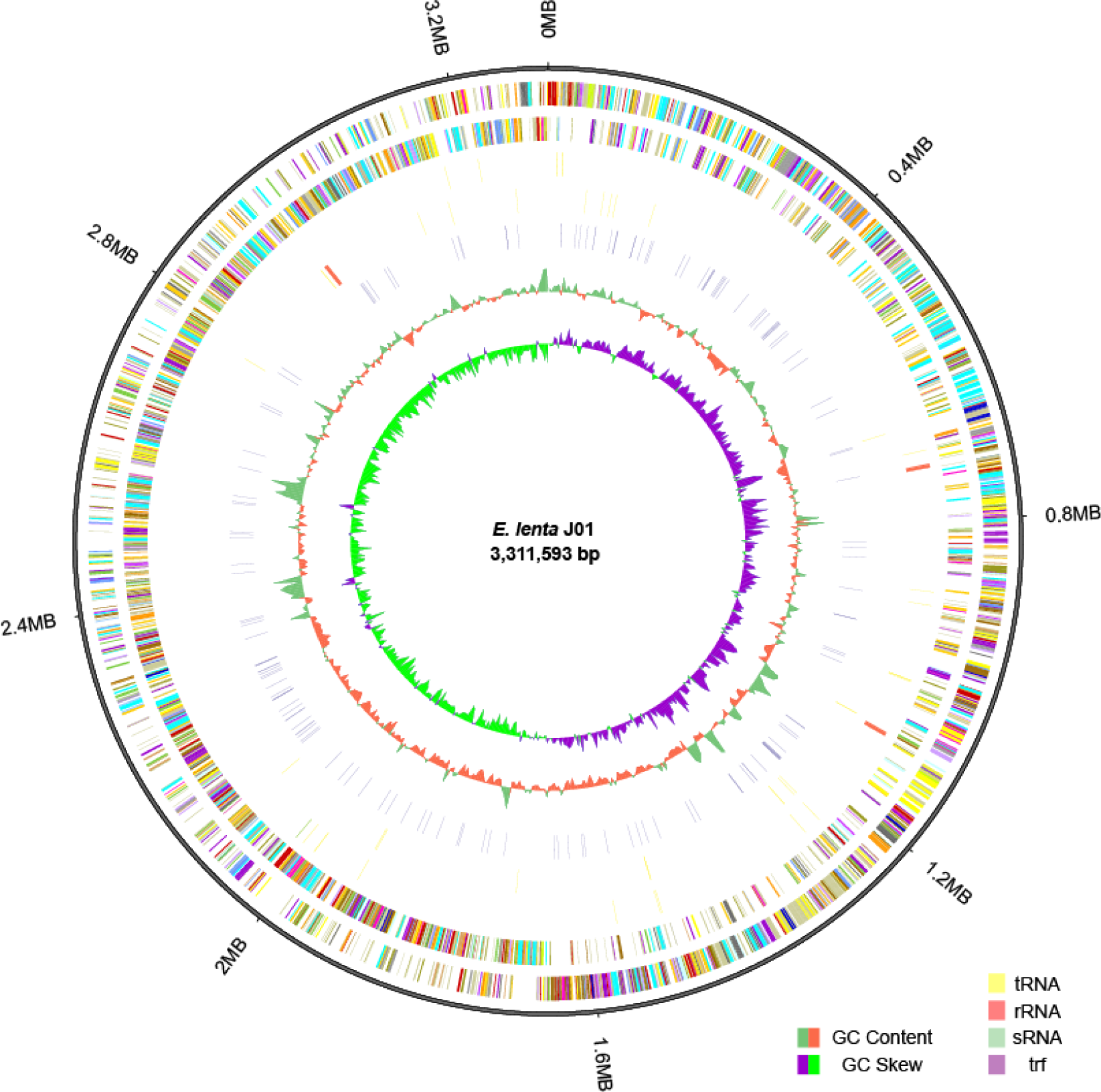
Genomic information of the *E. enta* J01 strain that exhibits efficient RSV-metabolizing ability. From outside to inside: circle 1, Genome Size; circle 2, Forward Strand Gene, colored according to cluster of orthologous groups (COG) classification; circle 3, Reverse Strand Gene, colored according to cluster of orthologous groups (COG) classification; circle 4, Forward Strand ncRNA; circle 5, Reverse Strand ncRNA; circle 6, repeat; circle 7, GC content; circle 8, GC-SKEW.

**Fig. S4.**
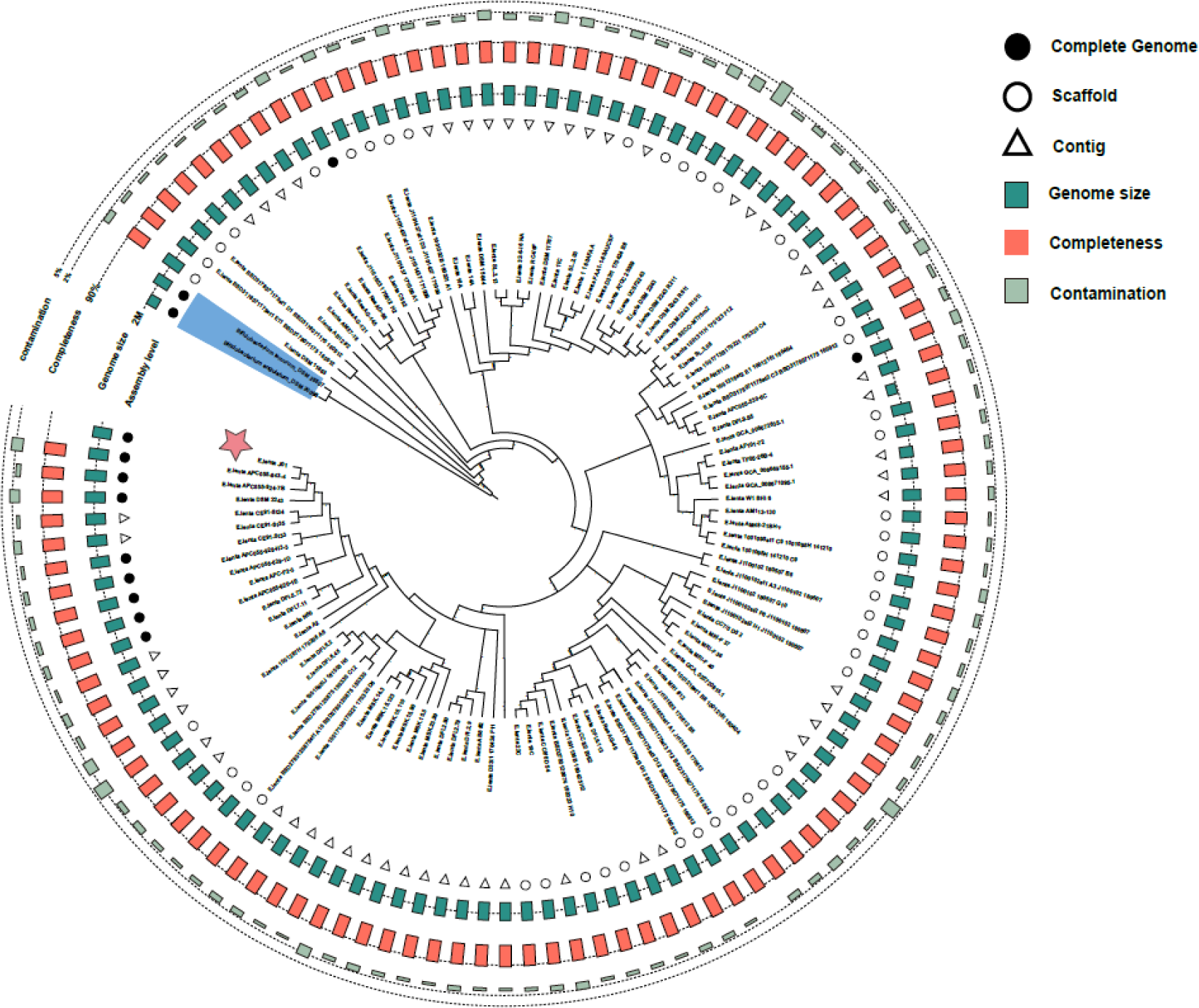
Phylogenetic analysis of *E. lenta* strains. From outside to inside: circle 1, contamination of genome; circle 2, completeness; circle 3, Genome size of strains involved in building phylogenetic trees; circle 4, The complete genome, scaffold, contig of the *E. lenta* strains involved in the construction of phylogenetic trees. In addition, the size of the column represents the level of the corresponding value.

**Fig. S5.**
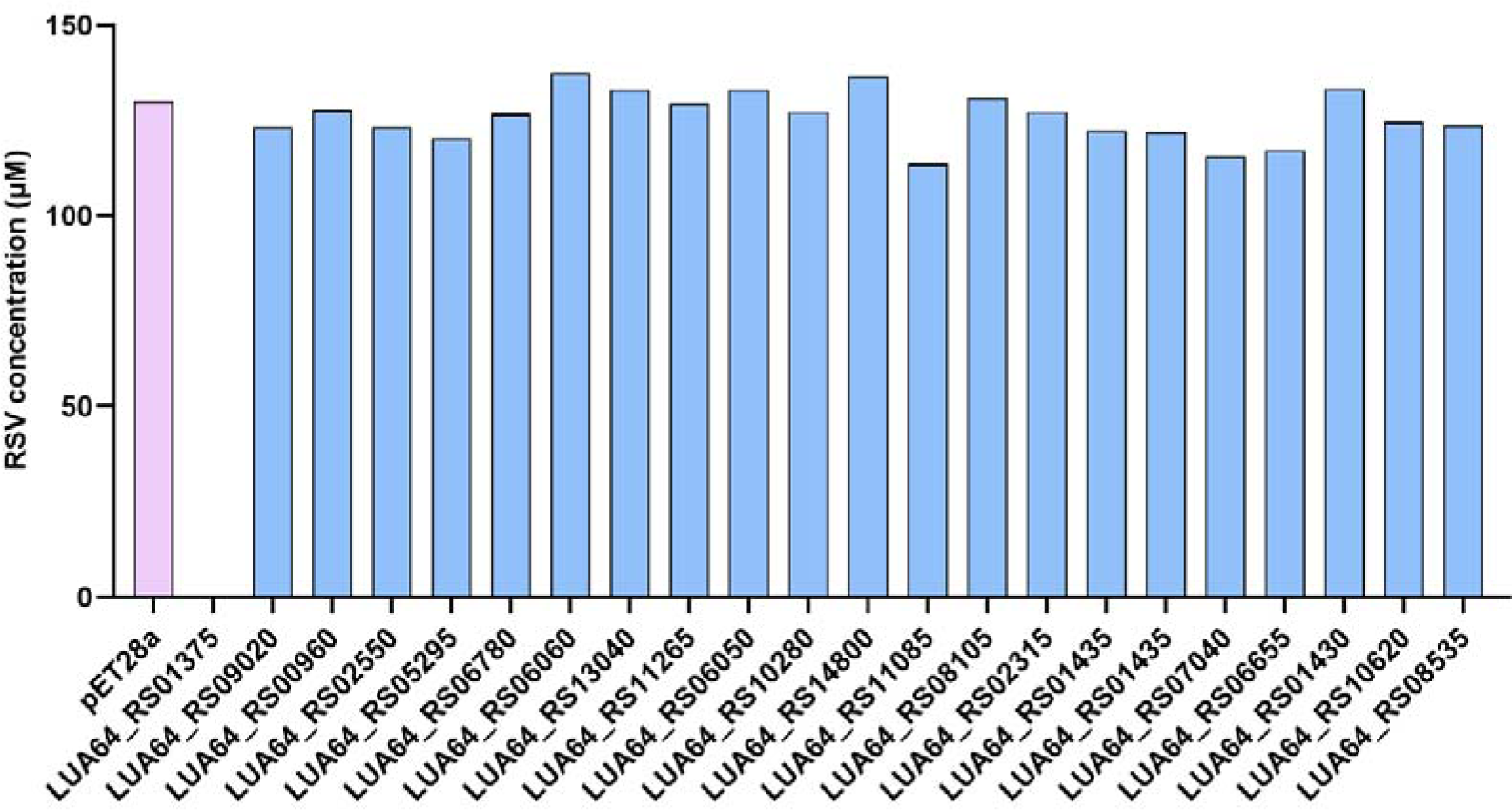
Screening for potential resveratrol reductases (RERs) The *E. coli* BL21 strains that expressed the predicted RER-encoding genes were cultivated with resveratrol. The consumption of resveratrol by the engineered *E. coli* BL21 strains was determined by measuring the residual resveratrol in the medium.

**Fig. S6.**
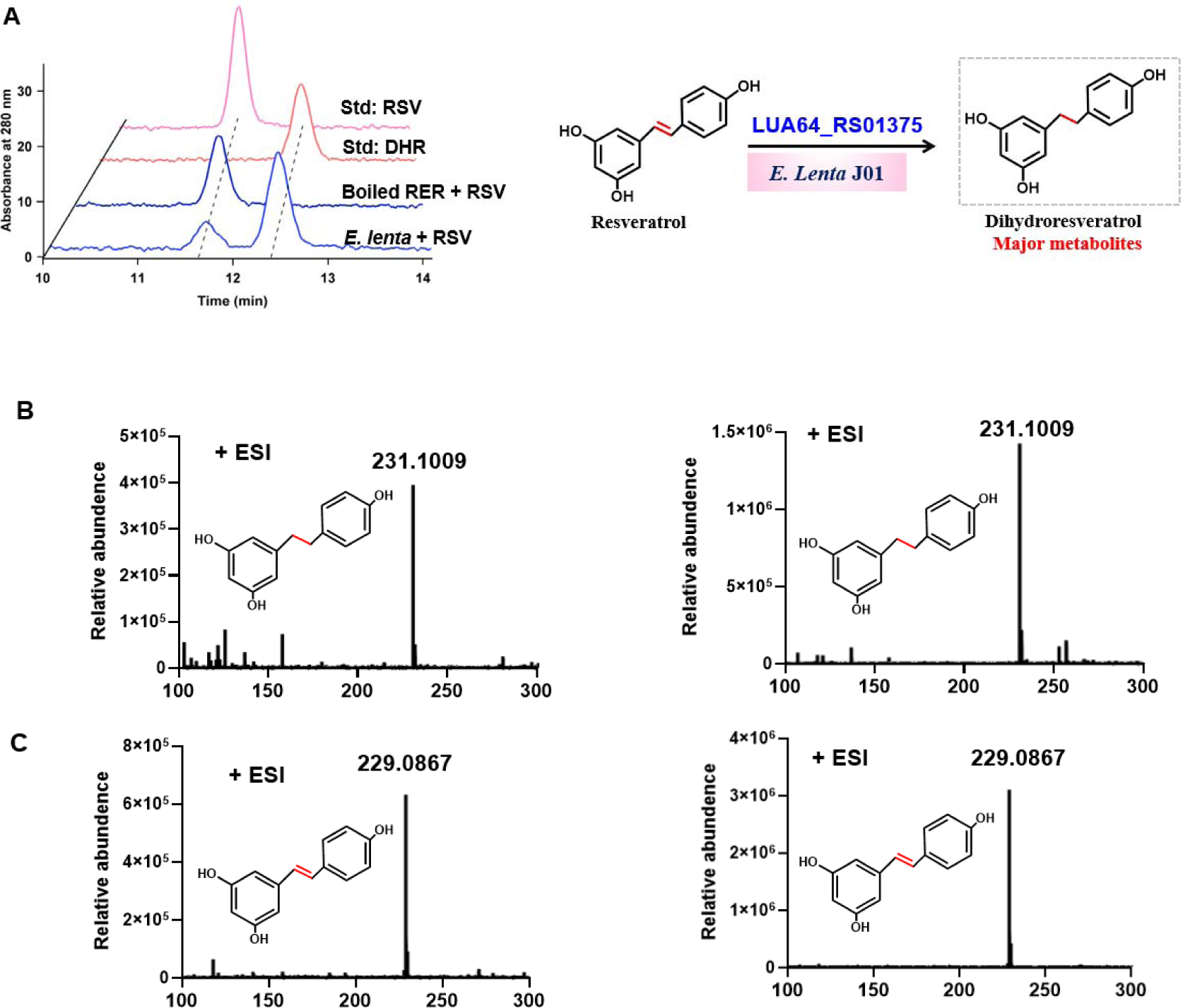
Mass spectrometry identification of RER-mediated transformation of RSV to DHR. **A**, Confirmation of the RER-catalyzed conversion of RSV to DHR. **B**, Tandem mass spectrometry analysis of the HPLC fractions corresponding to the main product formed by incubation of RER with RSV in vitro, and comparison with that of the DHR standard. **C**, Tandem mass spectrometry analysis of the HPLC fractions corresponding to the main product formed by incubation of RER with DHR in vitro, and comparison with that of the RSV standard.

**Fig. S7.**
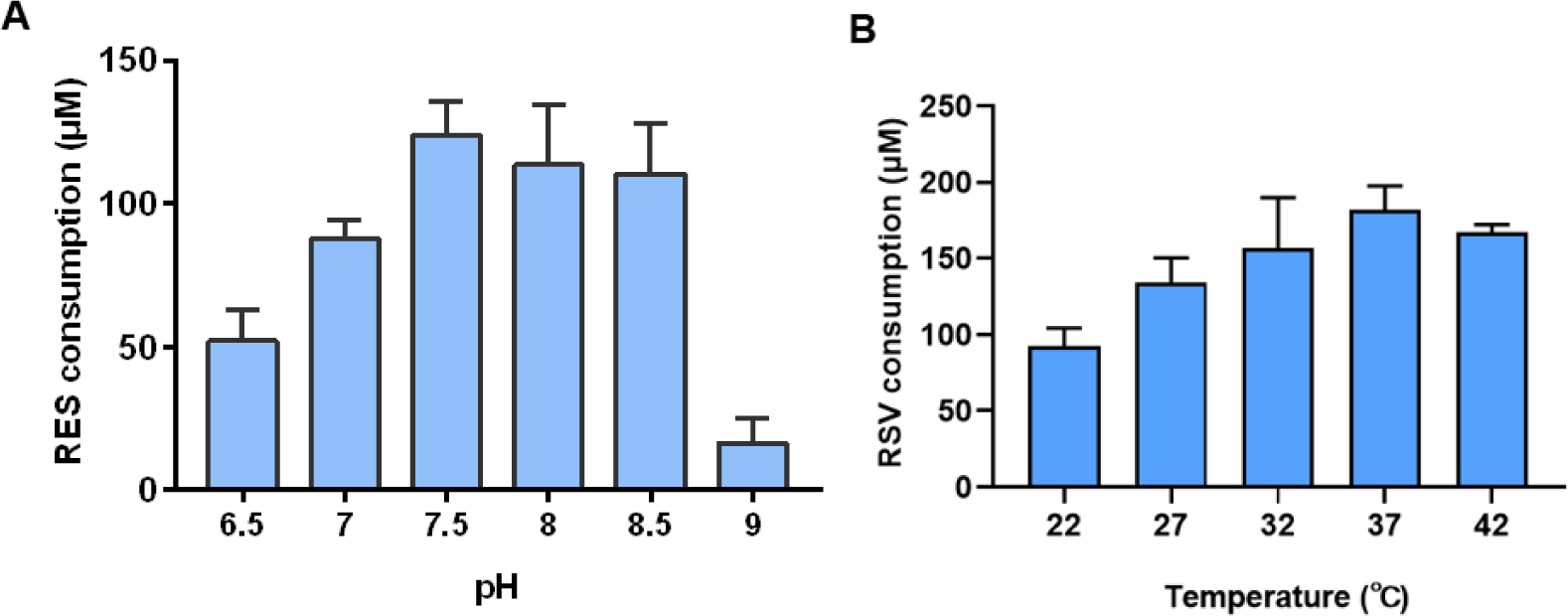
Determination of the optimum pH value and reaction temperature for steady-state kinetic analysis of RER using resveratrol as the substrate. **a** Determination of the optimum pH value. **b** Determination of the optimum reaction temperature. Data are represented as the mean ± s.d. (*n* = 3). Error bars show s.d.

**Fig. S8.**
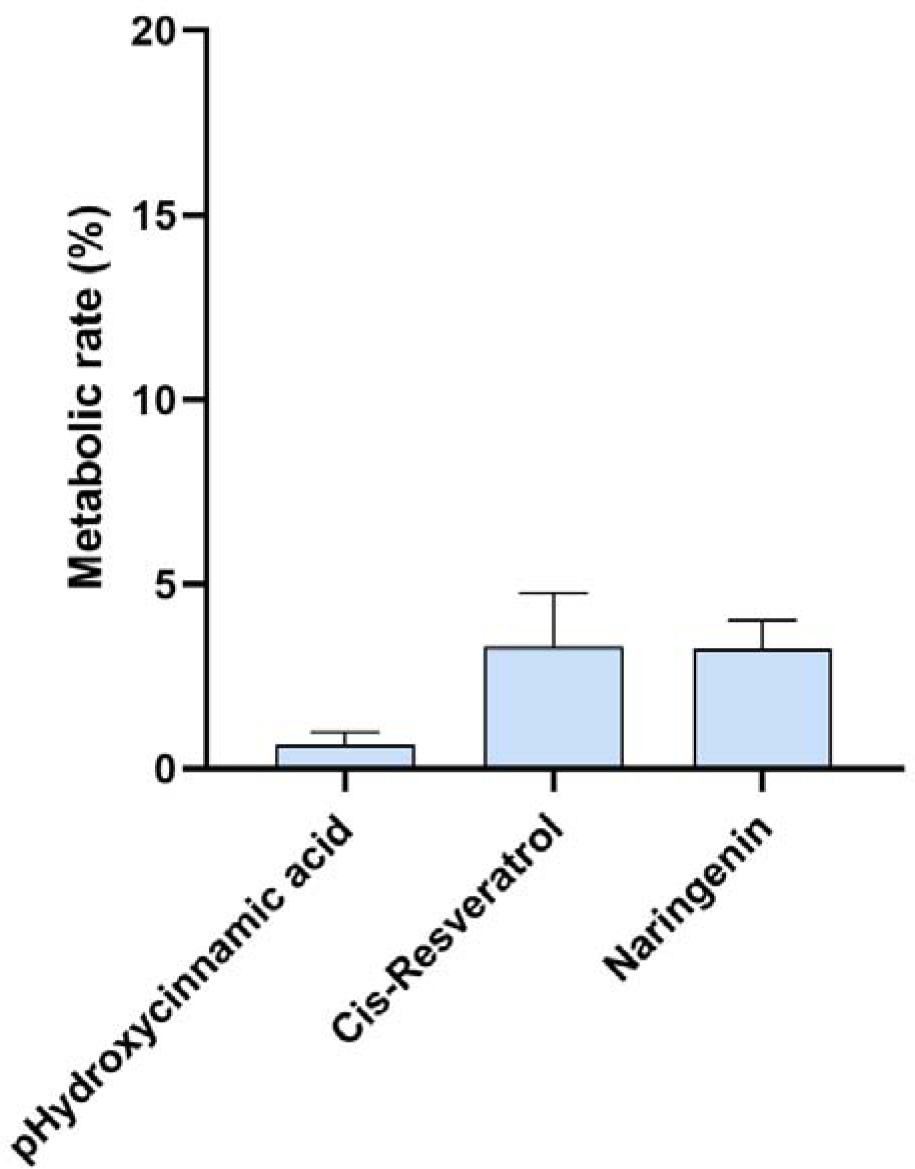
A screening of the substrate spectrum of RER. Initial turnover rates of the purified RER protein towards different substrates (100 μM) were determined. Data are represented as mean ± s.d. (*n* = 3).

**Fig. S9.**
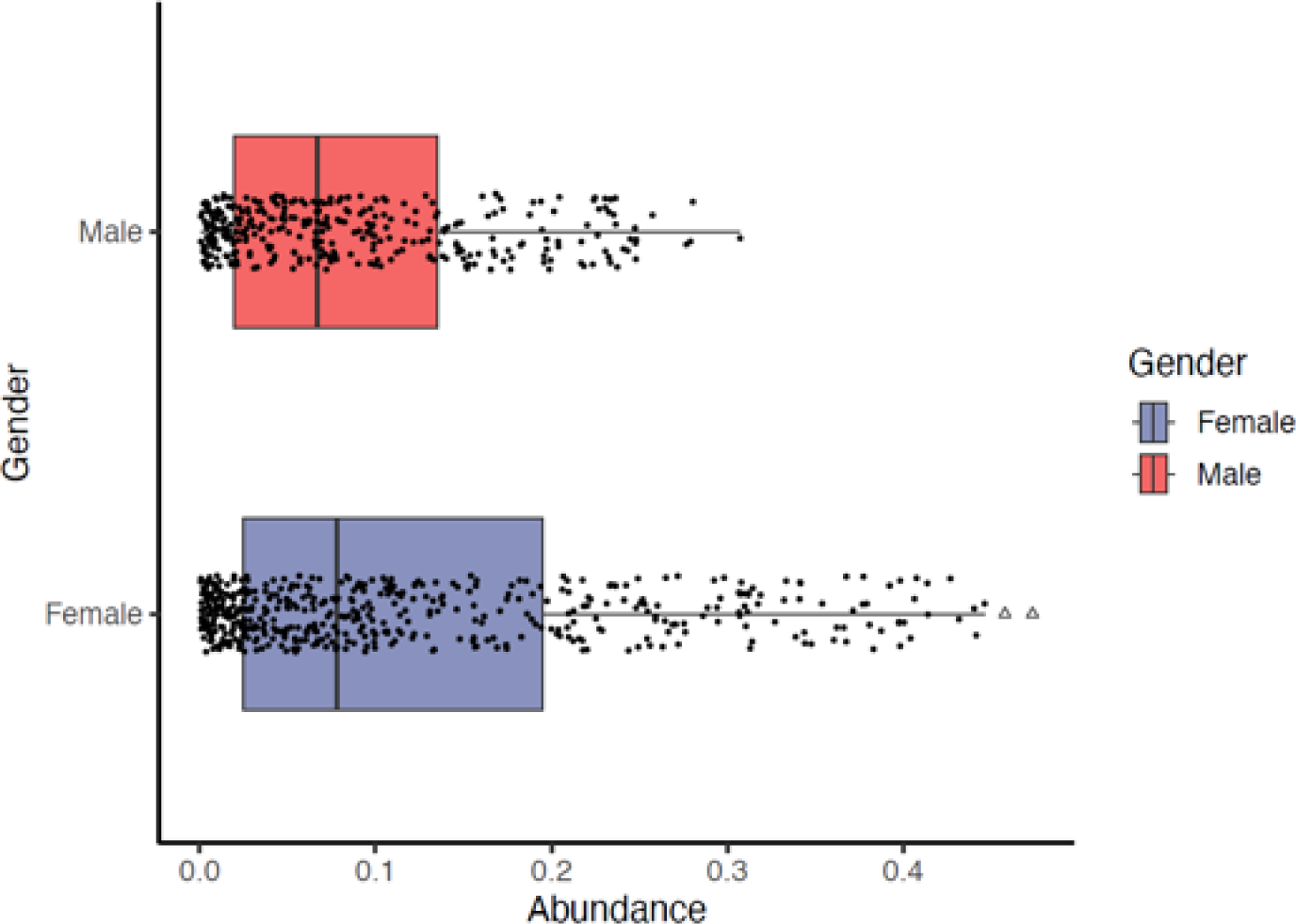
Abundance analysis of *rer*^+^ bacteria in male and female samples. The horizontal axis indicates the *rer* gene abundance, and the vertical axis indicates gender.

